# Phosphorylation of SAMHD1 Thr592 increases C-terminal domain dynamics, tetramer dissociation, and ssDNA binding kinetics

**DOI:** 10.1101/2022.04.05.486416

**Authors:** Benjamin Orris, Kevin W. Huynh, Mark Ammirati, Seungil Han, Ben Bolaños, Jason Carmody, Matthew D. Petroski, Benedikt Bosbach, David J. Shields, James T. Stivers

## Abstract

SAM and HD domain containing deoxynucleoside triphosphate triphosphohydrolase 1 (SAMHD1) is driven into its activated tetramer form by binding of GTP activator and dNTP activators/substrates. In addition, the inactive monomeric and dimeric forms of the enzyme bind to single-stranded (ss) nucleic acids. During DNA replication SAMHD1 can be phosphorylated by CDK1 and CDK2 at its C-terminal threonine 592 (pSAMHD1), enabling the enzyme to localize to stalled replication forks (RFs) and promote their restart. Since localization of a potent dNTPase at stalled RFs is not harmonious with DNA replication, we used a series of kinetic and thermodynamic measurements to explore a hypothesis where the combined effects of T592 phosphorylation and ssDNA binding serves as a dual switch to turn-off SAMHD1 dNTPase activity. We report that phosphorylation has only a small effect on the dNTPase activity and ssDNA binding affinity of SAMHD1. However, perturbation of the native T592 by phosphorylation decreased the thermal stability of tetrameric SAMHD1 and accelerated tetramer dissociation in the absence and presence of ssDNA (~15-fold). In addition, we found that ssDNA binds competitively with GTP to the A1 site. A full-length SAMHD1 cryo-EM structure revealed substantial baseline dynamics in the C-terminal domain (which contains T592) which may be modulated by phosphorylation. We propose that T592 phosphorylation increases tetramer dynamics and allows invasion of ssDNA into the A1 site and the previously characterized DNA binding surface at the dimer-dimer interface. These features are consistent with rapid and regiospecific inactivation of pSAMHD1 dNTPase at RFs or other sites of free ssDNA in cells.

## INTRODUCTION

SAM and HD domain containing deoxynucleoside triphosphate triphosphohydrolase 1 (SAMHD1) is the only known human dNTPase that catalyzes the breakdown of dNTPs into deoxynucleosides and inorganic triphosphate. The same activity also counteracts the effective cellular concentrations of many nucleoside-analog chemotherapeutics (1). This SAMHD1 activity is regulated through ordered binding of nucleotide activators to two allosteric sites on each monomer subunit of the enzyme, where GTP (or dGTP) binds to allosteric site one (A1) and any dNTP can bind to allosteric site two (A2) (2–4). Binding of eight activating nucleotides assembles SAMHD1 into a stable homotetramer capable of depleting dNTPs from nondividing or terminally differentiated cells, which is considered an innate immunity mechanism to inhibit replication of both DNA viruses and retroviruses in such cells (5–7). Notably, several viruses that infect humans encode proteins to counteract SAMHD1 by post-translational modification or targeted degradation via the polyubiquitinylation pathway (8–10). In addition, SAMHD1 has also been shown to bind single-stranded (ss) DNA and RNA at the dimer-dimer interface of the tetramer, which is strongly antagonistic to the dNTPase activity (11, 12). The cellular function of nucleic acid binding is not known, but such activity is consistent with the roles of the enzyme in double-strand break repair and restarting stalled replication forks (13–15).

SAMHD1 also has viral restriction and cellular DNA repair functions that do not involve dNTP hydrolysis but are correlated with phosphorylation of threonine 592 located in the largely disordered C-terminal domain (CtD) (16–19). In dividing cells, phosphorylation by CDK2 in early S phase is required for SAMHD1 to localize to stalled replication forks where it interacts with the MRE11 nuclease (20, 21), which promotes the degradation of nascent ssDNA and prevents ssDNA from spilling into the cytosol and activating the cGAS-STING pathway for interferon signaling (13, 22). In nondividing G0 macrophages—which are highly restrictive to viral infection—SAMHD1 is highly expressed, unphosphorylated, and dNTPase active (23, 24). However, macrophages also exist in a G1 state where SAMHD1 is phosphorylated at Thr592 by CDK1 (pSAMHD1), which is associated with permissiveness to viral infection, even though its dNTPase activity persists in the phosphorylated state (23, 25). Overall, data indicates that phosphorylation at Thr592 (pSAMHD1) serves to regulate SAMHD1 activity in a cell cycle dependent manner (16).

Given the regulatory importance of Thr592 phosphorylation, the effects of this post-translational modification have been the subject of biochemical, mutational, structural, and cell-based studies (16, 17, 26, 27). This work has generally found that phosphomimetic mutation of Thr592 (T592D or T592E), or in vitro kinase phosphorylation, produces relatively small effects on the dNTPase activity (2 to 4-fold)(2, 17, 18). One exception is that pSAMHD1 has a reduced dNTPase rate when dCTP is used as the substrate. The magnitude of this reduction has been reported to be in the range 3 to 10-fold (2, 27, 28). Several studies have also reported that phosphomimetic mutations destabilize the tetramer (17, 27, 28), which is supported by structural studies indicating that the mutants have an increased exposure of their activator nucleotide binding sites and a disruption of the dimer-dimer interface (27). These same phosphomimetic mutations have been shown to phenocopy the effects of phosphorylation in cells and *in vitro* biochemical measurements (14, 17, 21). Despite the importance of phosphorylation in regulating the cellular activity of SAMHD1 there is no established mechanism for how phosphoregulation occurs.

A paradox concerning the function of SAMHD1 is why this potent dNTPase is recruited to stalled replication forks where the requirement for dNTPs is high. An attractive resolution to this paradox is phosphorylation-induced inactivation of SAMHD1 dNTPase, but there is no evidence from *in vitro* or cell-based studies to support this mechanism. Here we explore another possible role where phosphorylation increases the dynamic motions of the SAMHD1 tetramer and allows ssDNA to invade the tetramer interface and abrogate dNTPase activity. This mechanism has the virtues of (i) providing regiospecific ablation of SAMHD1 dNTPase activity at cellular sites where the concentration of ssDNA is high, and (ii) preserving the bulk SAMHD1 dNTPase activity, which works in concert with ribonucleotide reductase to maintain balanced dNTP pools required for faithful DNA replication. We report that modification of Thr592 by phosphomimetic mutation, phosphorylation or deletion perturbs the A1 activator site and increases tetramer breathing dynamics. The increased dynamics allow invasion of ssDNA into the A1 site, leading to GTP release and further access to the previously characterized extended DNA binding site at the dimer-dimer interface. Thus, phosphorylation promotes rapid and regiospecific inactivation of pSAMHD1 dNTPase when ssDNA is present.

## METHODS

### Chemicals

2’-Deoxythymidine-5’-triphosphate (dTTP) and 2’-deoxyguanosine-5’-triphosphate (dGTP) were obtained from Promega. 2-^14^C labeled 2’-deoxythymidine-5’-triphosphate (2-^14^C-dTTP) was obtained from Moravek biochemicals. 2’-deoxyadenosine-5’-triphosphate (dATP) was obtained from New England Biolabs. Guanosine-5’-triphosphate (GTP) was obtained from Thermo Scientific. 2’-deoxythymidine-5’-[α-thio]-triphosphate (dTTPαS) and 2’-deoxyguanosine-5’-[α-thio]-triphosphate (dGTPαS) were obtained from Jena Biosciences. N-methylanthraniloyl guanosine-5’-triphosphate (mant-GTP) was obtained from AnaSpec. Glutaraldehyde was purchased from Sigma-Aldrich. C18 reversed-phase thin layer chromatography (TLC) plates were obtained from Macherey-Nagel. PEI cellulose TLC plates were obtained from EMD Millipore.

### DNA oligonucleotides and DNA sequencing

All DNA oligonucleotides were synthesized by Integrated DNA Technologies (IDT) and purified via polyacrylamide gel electrophoresis or HPLC. Sequences are listed **in Supplementary Table 1**.

### Protein expression and purification

Human SAMHD1 wild-type, T592E, Δ583-626, or Δ600-626 harbored in a pET19b plasmid as a PreScission protease cleavable 10xHis fusion construct was expressed in chemically competent BL21(DE3) E. coli cells (Agilent). An overnight starter culture of cells was grown in 2xYT supplemented with carbenicillin (50 μg/L) and subsequently inoculated 1:100 v/v in 2xYT media (shaker settings: 220 rpm, 37 °C) until an OD600 of 0.7 was reached, then cold-shocked for 30 minutes in an ice bath, induced with 1 mM isopropyl β-D-1-thiogalactopyranoside (ThermoFisher), and the cells were incubated for 20 hours (shaker settings: 180 rpm, 22 °C). Cells were harvested by centrifugation (10,000 x g) and stored at −80 °C.

Cell pellets were thawed and resuspended in lysis buffer (50 mM HEPES—pH 7.5, 300 mM KCl, 4 mM MgCl_2_, 0.5 mM TCEP, 25 mM imidazole, and 10% glycerol) comprising one tablet of protease inhibitor cocktail (Pierce), DNase I (Roche), 1 mg RNAse A (Alfa-Aesar), and 5 mg lysozyme (*Amresco*) per 50 mL of buffer. The resuspension was passed two times through a LM10 microfluidizer (Microfluidics) and centrifuged at 40,000 x g to produce a clarified lysate. The lysate was loaded onto a charged, pre-equilibrated 10 mL nickel column (HisPur Ni-NTA resin from ThermoFisher). The loaded column was given stringent, incremental washes with 30, 40, and 50 mM imidazole. After the UV trace returned to baseline, SAMHD1 was eluted from the column using 300 mM imidazole.

SAMHD1 isolates were dialyzed against 4 L of general buffer (50 mM HEPES—pH 7.5, 300 mM KCl, 4 mM MgCl_2_, 0.5 mM TCEP, and 10% glycerol) overnight with 1 mg GST-tagged PreScission protease added to remove the imidazole and 10xHis tag. The following day, the isolate was gently stirred at 4 °C with 2 mL glutathione-agarose resin, then centrifuged to pellet the resin and removed GST-PreScission protease. The SAMHD1 solution was concentrated to ~7 mg/ml and injected in 80 mg at a time onto a Cytiva Superdex 200 pg HiLoad 26/600 size exclusion column as a final purification step. The resulting SAMHD1 isolate was concentrated to 8-9 mg/ml and flash frozen in liquid nitrogen in 50 μL aliquots.

### In vitro enzymatic phosphorylation of SAMHD1

SAMHD1 (45 μM final concentration) and CDK2/Cyclin E1 (9 nM final concentration) were diluted in reaction buffer (40 mM HEPES pH 7.5, 10 mM MgCl_2_, 1 mM DTT, 0.005% Tween-20, 10% glycerol, 375 μM ATP) and incubated at room temperature for 1 hour with gentle mixing every 10 minutes. To assess the phosphorylation status of reacted SAMHD1, both reacted and unreacted SAMHD1 were characterized via mass spectrometry (Waters Acquity H-class UPLC system/Xevo G2-XS TOF) where reacted SAMHD1 had a mass shift consistent with the addition of a single phosphate group (**Supplementary Fig. S1a, b**).

To identify site(s) of phosphorylation, reacted SAMHD1 (6 mg) was denatured with 1:2 (v/v) of 3.2 M guanidine hydrochloride/1.6% formic acid prior to inline digest across a pepsin/protease XIII column (NovaBioAssays) and Nepenthesin 1 column (Phenomenex) using a 10-minute 8-38% acetonitrile gradient (200 ml/min) and subsequent MS2 by HCD on a Fusion Lumos mass spectrometer (ThermoFisher Scientific). Phosphorylated residues (serine, threonine, and tyrosine) were Sequest searched in Proteome Discoverer 2.2.0 software against the SAMHD1 fasta sequence (2 ppm mass error 0.05 Da fragment). The ptmRS algorithm was applied to provide further confidence in the assignment of phosphorylation. A single site, T592, was found to be phosphorylated (**Supplementary Fig. S1c**).

### Steady-state kinetic measurements and analysis

Standard reaction conditions for steady-state kinetic measurements were 0.01–5 mM GTP (activator), 0.1–5 mM dTTP (substrate), 10 nCi [2-^14^C]-dTTP, 50 mM HEPES - pH 7.5, 50 mM KCl, 5 mM MgCl_2_, and 0.5 mM TCEP in a 20 μL total reaction volume at 22 °C. For each GTP/dTTP concentration combination, thee replicate reactions carried out with SAMHD1 as the initiator. One-microliter fractions were withdrawn at regular time intervals and quenched by spotting onto a C18-reversed phase TLC plate. TLC plates were developed in 50 mM KH_2_PO_4_ (pH 4.0) to separate the substrate dTTP from the product dT. Developed TLC plates were exposed on a GE storage phosphor screen overnight and scanned on a Typhoon Imager (GE Healthcare). Substrate and product signal was quantified with GelBandFitter using a Gaussian fitting algorithm. The amount of product formed at each time point in each replicate was calculated using eq 1:

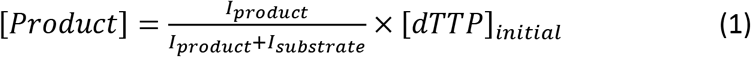

where I_product_ is the signal intensity of the dT product peak, I_substrate_ is the signal intensity of the dTTP substrate peak, and [dTTP]_initial_ is the initial concentration of dTTP substrate in the reaction. Initial rates of product formation were obtained from plots of [dT] vs. time, with rates corresponding to slope, and error corresponding to standard error of the slope as determined by linear regression analysis. Reaction rates (μM/minute) were plotted vs. dTTP substrate concentration using five fixed concentrations of GTP activator.

For GTP activation with dTTP as the substrate, the kinetic parameters were determined by fitting to an ordered essential activation mechanism (eqs 2-4)(4), where [dTTP] and [GTP] are the free nucleotide concentrations, *V*_max_^app,dTTP^ is the apparent maximal velocity for dTTP hydrolysis at a given activator concentration, *K*_m_^app,dTTP^ is the apparent Michaelis constant for dTTP at a given concentration of GTP activator, *K*_act_^GTP^ is the activation constant for GTP, *K*_m_^dTTP^ is the *K_m_* for dTTP in the absence of [GTP] and α is a unitless constant indicating the degree to which *K*_m_^app,dTTP^ is decreased at saturating [GTP]. The reported *k_cat_* values were calculated from the equation *k*_cat_ = *V*_max_^dTTP^/[SAMHD1 monomers], where *V*_max_^dTTP^ is the value of *V*_max_^app,dTTP^ at saturating [GTP].

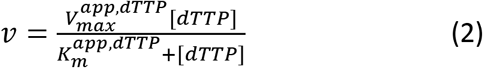

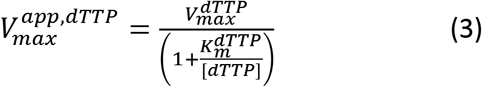

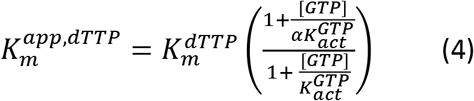

### Fluorescence anisotropy measurements of ssDNA binding

The fluorescence anisotropy measurements were carried out in quartz microcuvettes in a FluoroMax 3 Spectrofluorometer (Horiba Scientific) maintained at 25°C. Binding reactions were prepared in binding buffer (50 mM HEPES [pH 7.5], 50 mM KCl, 1 mM EDTA, .5 mM TCEP) with 50 nM of 5’ FAM-labeled oligonucleotide. Anisotropy of the FAM fluorophore was measured (excitation and emission wavelengths of 493 and 517 nm; 0.5 s integration time) as increasing concentrations of SAMHD1 were added. The resulting anisotropy vs. total [SAMHD1] curves were fit to a quadratic binding equation (eq 6):

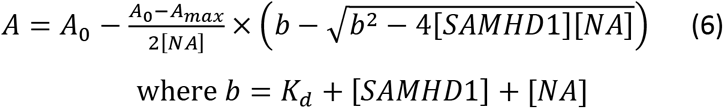

where A is the observed anisotropy, A_0_ is the initial anisotropy value, Amax is the maximal anisotropy value at saturation, [NA] is the total nucleic acid concentration, and [SAMHD1] is the total SAMHD1 concentration. The same anisotropy assay was used to measure competitive displacement of ssDNA by GTP using [GTP] in the range 0.01 to 1 mM. The apparent dissociation constants for DNA obtained at each [GTP] (*K*_D_^app, DNA^) were fitted to the competitive binding equation *K*_D_^app, DNA^ = *K*_D_^DNA^ + [GTP] x (*K*_D_^DNA^/ *K*_D_^GTP^).

### Mant-GTP fluorescence measurements

Mant-GTP fluorescence intensity measurements were carried out in quartz microcuvettes in a FluoroMax 3 Spectrofluorometer (Horiba Scientific) maintained at 25°C. Binding reactions were prepared in binding buffer (50 mM HEPES [pH 7.5], 50 mM KCl, 1 mM EDTA, 0.5 mM TCEP) with 500 nM mant-GTP. Emission intensity of the mant fluorophore was measured (excitation and emission wavelengths of 335 and 440 nm; 0.1 s integration time) as increasing concentrations of SAMHD1 were added. The raw fluorescence data was corrected for dilution, inner-filter effects, and photobleaching using the following formula:

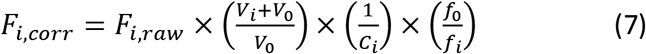

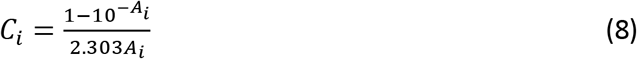

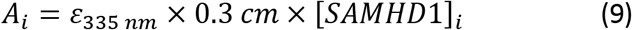

Where F_i,corr_ is the corrected fluorescence at the i^th^ point in the titration, F_i,raw_ is the raw fluorescence at the i^th^ point in the titration, *V*_i_ is the total volume of SAMHD1 solution added up to the i^th^ point in the titration, *V*_0_ is the starting volume of the titration, A_i_ is the total absorbance of SAMHD1 at the 335 nm excitation wavelength of mant-GTP at the i^th^ point in the titration, f_o_ is the starting fluorescence of mant-GTP in a mock titration with no SAMHD1 being added, and f_i_ is the fluorescence of mant-GTP at the i^th^ point in the mock titration. Following correction, the data was normalized to percent fluorescence increase and fit to a quadratic binding equation:

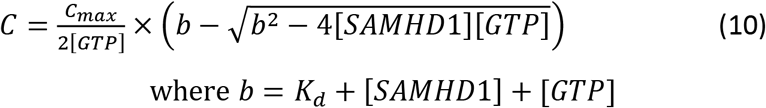

where C is the observed percent fluorescence increase, C_max_ is the maximal percent fluorescence increase at saturation, [GTP] is the total concentration of mant-GTP, and [SAMHD1] is the total SAMHD1 concentration.

### Thermal melt measurements

Thermal melt measurements were using a GloMelt Thermal Shift protein stability kit from Biotium on a Qiagen Rotor-gene Q qPCR instrument. Solutions of 50 mM HEPES – pH 7.5, 50 mM KCl, 5 mM MgCl_2_, 0.5 mM TCEP, 3 uM SAMHD1, 1X GloMelt dye, and 0/2 mM dGTPαS were subjected to a temperature gradient from 25 to 85 °C at 0.5 °C intervals. Each step in the gradient was held for three seconds and the fluorescence of the GloMelt dye was measured Rotor-gene Q green channel. Automatic gain optimization was performed all of the samples before execution of the temperature gradient with a signal limit of seven. Three replicates were recorded for each measured condition. Melt temperatures were calculated for each replicate were calculated using the program TSA-CRAFT (**Supplemental Table S2**)(29). To display differences in melt temperature between different SAMHD1 constructs, averaged traces were differentiated and normalized in Prism using 6^th^ order polynomial smoothing considering 5 neighbors on each side.

### Single-stranded DNA competition of dNTPase activity

Reaction conditions for ssDNA competition experiments were 10 μM GTP, 10 μM dTTP, 10 nCi [2-^14^C]-dTTP, 0/10/20/50 μM ssDNA90, 50 mM HEPES - pH 7.5, 50 mM KCl, 5 mM MgCl_2_, and 0.5 mM TCEP in a 20 μL total reaction volume at 22 °C. For each ssDNA90 concentration, thee replicate reactions were carried out with SAMHD1 as the initiator. One-microliter fractions were withdrawn at regular time intervals and quenched by spotting onto a PEI cellulose TLC plate. TLC plates were developed in 100 mM LiCl to separate the substrate dTTP from the product dT. Developed TLC plates were exposed, imaged, and quantified as previously described in “Steady-State Kinetic Analysis”, with the exception that quantification was performed using ImageJ. Each rate is the average initial rate of three replicate reactions with error bars representing standard error of the slope determined by linear regression analysis.

### Tetramer lifetime measurements

A 100 μL solution of 1 mM dGTP, 50 mM HEPES – pH 7.5, 50 mM KCl, 5 mM MgCl_2_, 0.5 mM TCEP, and 25 μM SAMHD1 was incubated for 30 seconds, then diluted 100X into a solution of 50 mM HEPES – pH 7.5, 50 mM KCl, 5 mM MgCl_2_, and 0.5 mM TCEP with no activating nucleotides. Three replicate reactions were done with each enzyme. Fractions were withdrawn at regular time intervals for 8 hours and crosslinked in 50 mM glutaraldehyde for 10 minutes. Crosslinking was halted by the addition of Tris – pH 7.5. Crosslinked fractions were loaded onto a 1.5 mm, 10-well, 4-12% acrylamide gradient Bis-Tris gel (Invitrogen) and ran at 200 V for 40 minutes to separate tetrameric, dimeric, and monomeric species. Gels were stained with Coomassie R250 dye. A PAGEruler pre-stained molecular weight ladder (Thermo) was used to identify which bands corresponded to the different oligomeric states of SAMHD1. Traces were fitted to a two-phase exponential decay model using GraphPad Prism.

### ssDNA binding kinetic measurements

A 20 μL solution of 1 mM dGTP, 50 mM HEPES – pH 7.5, 50 mM KCl, 5 mM MgCl_2_, 0.5 mM TCEP, and 25 μM SAMHD1 was incubated for 30 seconds, then diluted 100X into a solution of 50 mM HEPES – pH 7.5, 50 mM KCl, 1 mM MgCl_2_, 0.5 mM TCEP, and 50 nM 5’FAM-labeled ssDNA57 with no activating nucleotides. Five replicate reactions were done with each enzyme. Control reactions with no dGTP in the pre-dilution mixture were also carried out. Anisotropy of the FAM fluorophore was recorded at 15 second intervals for 10 minutes after dilution. Traces were fitted to one-phase association model using GraphPad Prism.

### Course-grained normal mode analysis

A structural model for the SAMHD1 monomer was extracted from the tetramer structure (PDB: 6TXC) (30). Normal mode analysis was carried out on the monomer chain in Rstudio using the package Bio3D (31). Fluctuations and deformation energy were mapped on the SAMHD1 structure as b-factors and displayed in Pymol using the putty cartoon representation. The R-script file is provided in **Supplemental Methods.**

### T592V to T592E structure morph

A 40-frame video morph of SAMHD1 T592V (PDB: 4ZWE) to SAMDH1 T592E (PDB: 4ZWG) (27) as generated in the ChimeraX program using the morph function on extracted monomers from the tetrameric crystal structures (32). Alpha-carbon fluctuations between the structures were determined by aligning the structures in Pymol and calculating paired atom displacements using the colorbyrmsd function. Results were mapped on the structure as b-factors and exported for comparison with course-grained normal mode analysis results.

### Negative Stain EM

The SAMHD1-dGTPαS complex prepared for cryo-EM was diluted in buffer containing 50 mM HEPES pH 7.5, 0.3 M KCl, 0.5 mM TCEP, 2 mM MgCl_2_, and 1 mM dGTPαS, applied to glow discharged (Pelco EasiGlow, 0.19 mBar, −15 mA, 20 s) copper grids with a thin carbon film (CF300-Cu-UL, EMS), washed with water, and stained with 2% uranyl acetate. Grids were imaged at 50 kx (2.087 A/pix) using a 200 keV Tecnai TF20 transmission electron microscope (ThermoFisher Scientific) equipped with a OneView camera (Gatan). Grids of SAMHD1 prior to nucleotide addition were prepared in the same manner. Data was collected using Serial EM and processed with Relion 3.1.0 to observe 2D class averages of the nucleotide induced SAMHD1 tetramer (33)(**Supplemental Figure S2**).

### Preparation of SAMHD1-dGTPαS complex grids for cryo-EM

To form the SAMHD1 tetramer, purified protein was thawed and diluted to 0.6 mg/mL with buffer containing 50 mM HEPES pH 7.5, 0.3 M KCl, 0.5 mM TCEP, 2 mM MgCl_2_, and 1 mM dGTPαS. After incubation on ice for 30 min, the complex was diluted to 0.25 mg/mL, applied to plasma cleaned (ArO_2_) Quantifoil R1.2/1.3 200 mesh Au grids, and vitrified in liquid ethane (Vitrobot Mark IV ThermoFisher Scientific).

### Data acquisition

Dose fractionated movies were acquired using SerialEM from a Titan Krios G2 transmission electron microscope (ThermoFisher Scientific) operating at 300 keV equipped with a K2 Summit direct electron detector and Quantum LS energy filer (Gatan) (34). Three-thousand two-hundred and twenty-two movies were collected in super-resolution mode with a pixel size of 0.42 Å/pixel, using a 100μm objective aperture, 20 ev slit, and a dose rate of 8.16 e-/Å^2^/s over a defocus range of −0.6μm to −3.2μm with 7.03 sec exposures for a total of 38 frames.

### Cryo-EM Data Processing

All data processing was performed in RELION 3.1.0 unless otherwise stated (33). Individual frames from each of the 3,222 raw movies were aligned using MotionCor2 v1.0.5 and CTF estimated using CTFFIND4 (35, 36). In preparation for RELION reference-based auto-picking, 2,064 particles were manually picked from the first 500 micrographs and classified into ten 2D class averages. Four 2D class averages of different views of the particle were selected for reference-based auto-picking, resulting in 1,030,155 picked particles.

Particles were extracted with a box size of 300 pixels, then binned by 2 to 150 pixels before 2D classification. The first round of 2D classification yielded 36 2D classes with 661,791 particles with either secondary structure features or high-resolution information. These particles were selected and unbinned for another round of 2D classification, yielding 495,583 in 29 class averages. An *ab initio* 3D initial model generated in RELION 3.1.0 was used for 3D classification. One 3D class average of 114,078 particles, with clear α-helical and β-strand features, was selected for 3D auto-refinement.

The first round of 3D auto-refinement yielded a 3.15Å resolution structure after postprocessing. Two iterations of CTF refinement and one iteration of particle polishing followed by 3D auto-refinement and postprocessing generated a final cryo-EM map of 2.89Å resolution, based on the FSC 0.143 criteria. The final cryo-EM map was flipped. Local resolution analysis was calculated using ResMap (37). The map was also sharpened using the *phenix.auto_sharpen* command in the Phenix software package version 3594 (38).

### Atomic modeling and model refinement

The structure of the human SAMHD1-GTP-dGTP complex (PDB: 4TNX) was fit into the final structure using UCSF Chimera (39). The atomic model was built in COOT (40). The ligand dGTPαS (PDB: T8T) was modeled into allosteric sites 1 and 2 and the substrate binding site. Phenix real space refinement was performed to further optimize the model. The overall cryo-EM data processing workflow is shown in **Supplemental Figure S3,** and the structural statistics are reported in **Supplementary Table S3**.

## RESULTS

### Steady-state dNTPase activity of SAMHD1, T592E and pSAMHD1

Since several conflicting reports exist on the dNTPase activity of pSAMHD1 and the T592E phosphomimetic form, and N-terminal SAM-domain deletion constructs of differing lengths have been employed by various groups, we performed a thorough steady-state kinetic analysis using full-length wild-type SAMHD1 (**Fig. 1a**) and its T592E variant (**Fig. 1b**). We employed [2-^14^C] labeled dTTP as the substrate and used a previously described reversed-phase TLC assay for resolving the dTTP substrate from the thymidine product (**Supplemental Fig. S4**). Reactions for SAMHD1 and T592E were performed using 0.5 μM [enzyme] and both GTP activator and substrate were varied in the concentration range 0.01 to 5 mM. The data were fit to eq 1-4 to obtain *k*_cat_, *K*_act_, ^GTP^, *K*_m_^dTTP^ and *α* for SAMHD1 and the T592E variant (**Fig. 1a and 1b**). Like a subset of the previous reports, our analysis revealed only small differences in the kinetic parameters between SAMHD1, T592E (*k*_cat_^dTTP,WT^ = 4.9 s^-1^, *k*_cat_^dTTP,T592E^ = 4.9 s^-1^, *K*_act_^GTP,WT^ = 194 μM, *K*_act_^GTP,T592E^ = 73 μM, *K*_m_^dTTP,WT^ = 4.3 mM, and K_m_^dTTP,T592E^ = 4.4 mM) (**Supplemental Table S4**). A secondary replot of *K*_m_,^app,dTTP^ versus [GTP] shows that SAMHD1 and T592E possess almost no affinity for dTTP in the absence of the activator GTP (**Fig. 1c**). This establishes that for all realistic concentrations of dTTP found in cells, GTP is an essential *K*_m_-type activator for both enzymes. Since pSAMHD1 was not as abundantly available as SAMHD1 and T592E, we performed a more limited kinetic analysis on pSAMHD1 using several concentrations of GTP and dTTP to ascertain whether its behavior was significantly different from T592E (**Fig. 1d)**. We observed only small differences between T592E and pSAMHD1, indicating that the phosphomimetic serves as a reasonable surrogate for pSAMHD1 in the context of steady-state kinetic measurements over a wide range of activator and substrate concentrations (**Fig. 1d**). Thus, these data establish that the function of phosphorylation is not to “switch off” the dTTPase activity. This conclusion extends to dGTP and dATP substrates based on several previous reports (2, 28), but dCTP appears to be a special case where the dNTPase activity is selectively reduced for T592E (2, 28).

**Figure 1.**
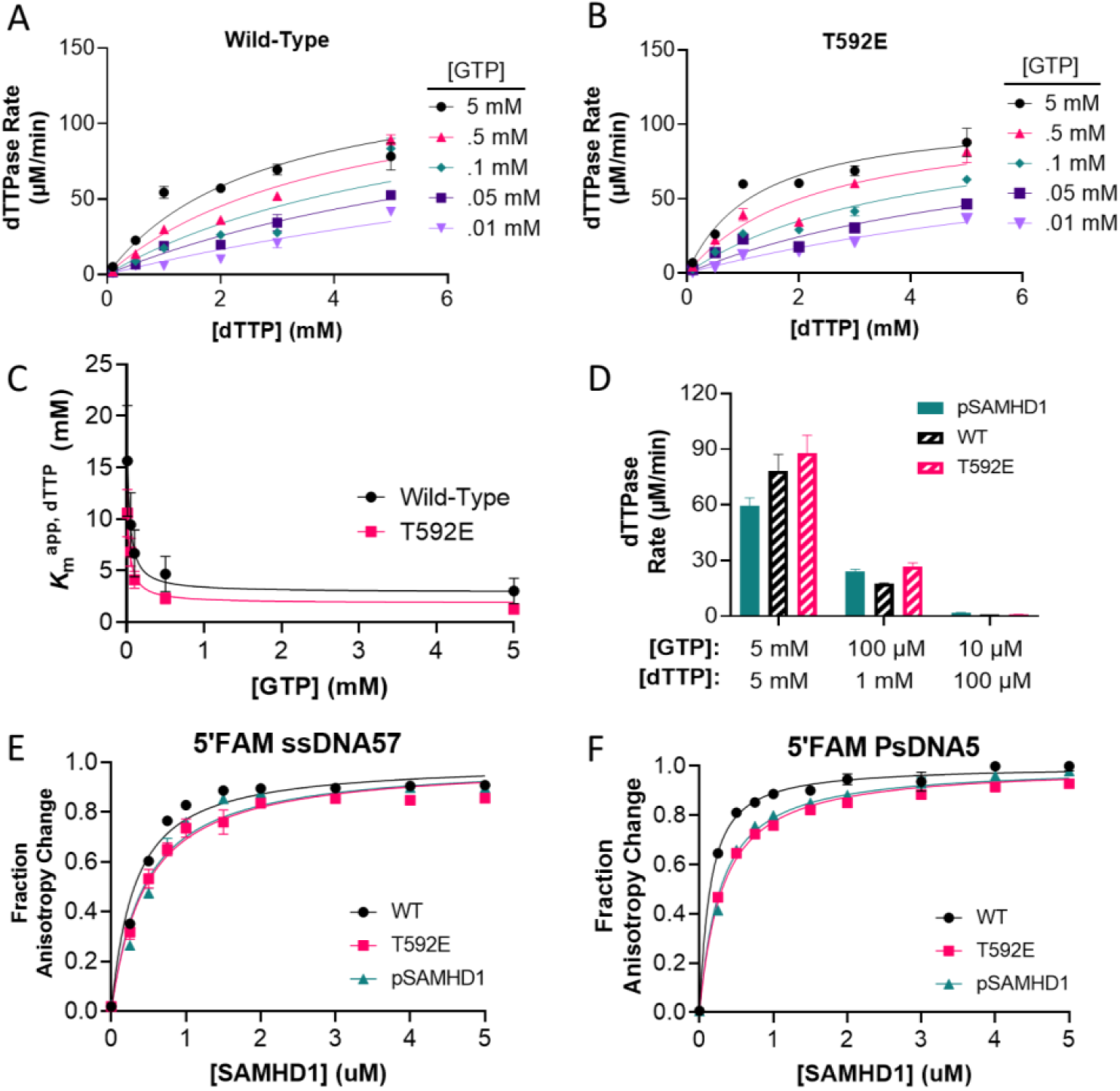
Both the T592E phosphomimetic mutation and phosphorylation have no significant impact on the dNTPase activity or DNA-binding affinity of SAMHD1. **(A)** Kinetic analysis of wild-type SAMHD1. SAMHD1 (0.5 μM) was incubated with varying concentrations of GTP and [2-^14^C]-dTTP. Curves represent least-squares regression fit to the Michaelis-Menten equation. **(B)** Kinetic analysis of SAMHD1 T592E under the same conditions as in (A). Error bars in (A) and (B) indicate standard error of reaction rate determined by the linear regression fit of the linear phase of three replicate reactions performed for each condition. **(C)** Secondary replot of apparent K_m_ with respect to dTTP substrate concentration as a function of GTP activator concentration. Error bars indicate standard error of the K_m_ values determined in (A) and (B). **(D)** dNTPase activity of pSAMHD1 compared with wild-type SAMHD1 and SAMHD1 T592E under select conditions from the kinetic analysis in (A) and (B). Error bars indicate standard error of the slope determined by the linear regression fit of the linear phase of three replicate reactions under each condition. **(E)** Binding of SAMHD1 wild-type T592E, and pSAMHD1 to 5’FAM-labeled ssDNA57 (50 nM). **(F)** Binding of SAMHD1 wild-type, T592E, and pSAMHD1 to 5’FAM-labeled PsDNA5 (50 nM). Error bars in (E) and (F) indicate the standard error of mean determined by three replicate titrations.

### ssDNA binding to SAMHD1, T592E and pSAMHD1

Since our model invokes preferential inactivation of pSAMHD1 in the presence of ssDNA, we compared the ssDNA binding affinity of SAMHD1, T592E and pSAMHD1 using two ssDNA ligands and a previously described fluorescence anisotropy assay (**Fig. 1 e, f).** Both ssDNA ligands contained a 5’-FAM label but differed in length and composition of the phosphate backbone linkages. The first DNA was a mixed sequence 57mer (ssDNA57) that has been used previously in DNA binding studies of SAMHD1 and contains normal phosphodiester linkages and a guanine rich 5’ sequence (5’-TGGAG-3’)(11, 12). The second DNA was a short 5mer containing racemic phosphorothioate substitutions (*) at its four phosphate linkages (5’FAM-dC*G*C*C*T) (FAM-PsDNA5). PsDNA5 was chosen based on a previous report that this construct binds SAMHD1 with high binding affinity, and a crystal structure reveals that the dG residue of PsDNA5 binds specifically in the A1 site like GTP, while the dC nucleotides dock across the A2 site (PDB: 6U6X)(41).

Anisotropy-based binding experiments were performed by adding small portions of a concentrated SAMHD1 solution to 50 nM [DNA] and recording the anisotropy increases of the FAM fluorophore after each addition. Using this approach, we found no notable differences in binding affinity of SAMHD1, T592E or pSAMHD1 for ssDNA57 or PsDNA5 (**Fig. 1e, f**). Consistent with a previous report, the PsDNA5 bound with similar affinity to ssDNA57 despite its shorter length: *K*_D_^PSDNA5^ = 119 nM; *K*_D_^ssDNA57^ = 286 nM. We conclude that neither phosphomimetic mutation nor phosphorylation at Thr592 perturbs binding of SAMHD1 monomers to long or short ssDNA constructs in the absence of nucleotides.

### GTP displaces bound ssDNA from SAMHD1 and its T592 variants

Based on the structural observation that PsDNA5 bound to the A1/A2 activator sites we investigated whether GTP and the non-hydrolysable substrate analogue dTTPαS impacted DNA binding to SAMHD1, T592E and pSAMHD1 (**Figure 2**). The three enzyme forms were titrated into solutions of 50 nM 5’ FAM-labeled ssDNA57 or PsDNA5 in the presence of increasing concentrations of GTP in the range 0.01 to 1 mM. The data for ssDNA57 are shown in **Figure 2a** and the data for PsDNA5 are shown in **Supplemental Figure S5**. For all three enzymes, the apparent *K_D_* values for ssDNA57 and PsDNA5 increased with a linear dependence on [GTP] (**Fig. 2b**), consistent with competitive binding of GTP with both short and long DNA ligands. The relative slopes for GTP competition in Figure 2d are inversely related to the respective binding affinities of the bound DNA as required by the equation for competitive binding (see Methods) (i.e., the tighter binding PsDNA5 requires higher concentrations of GTP for displacement). When both 1 mM dTTPαS and 1 mM GTP were added to SAMHD1 to induce tetramerization, there was also no detectable DNA binding (**Fig. 2a**) confirming that the tetramer is not the form of SAMHD1 competent for DNA binding. These results suggest that saturation of the A1 site using 1 mM GTP is sufficient to increase the mobility of the 5’ DNA end. The simplest interpretation of this result is that the DNA end is in proximity to the A1 site.

**Figure 2.**
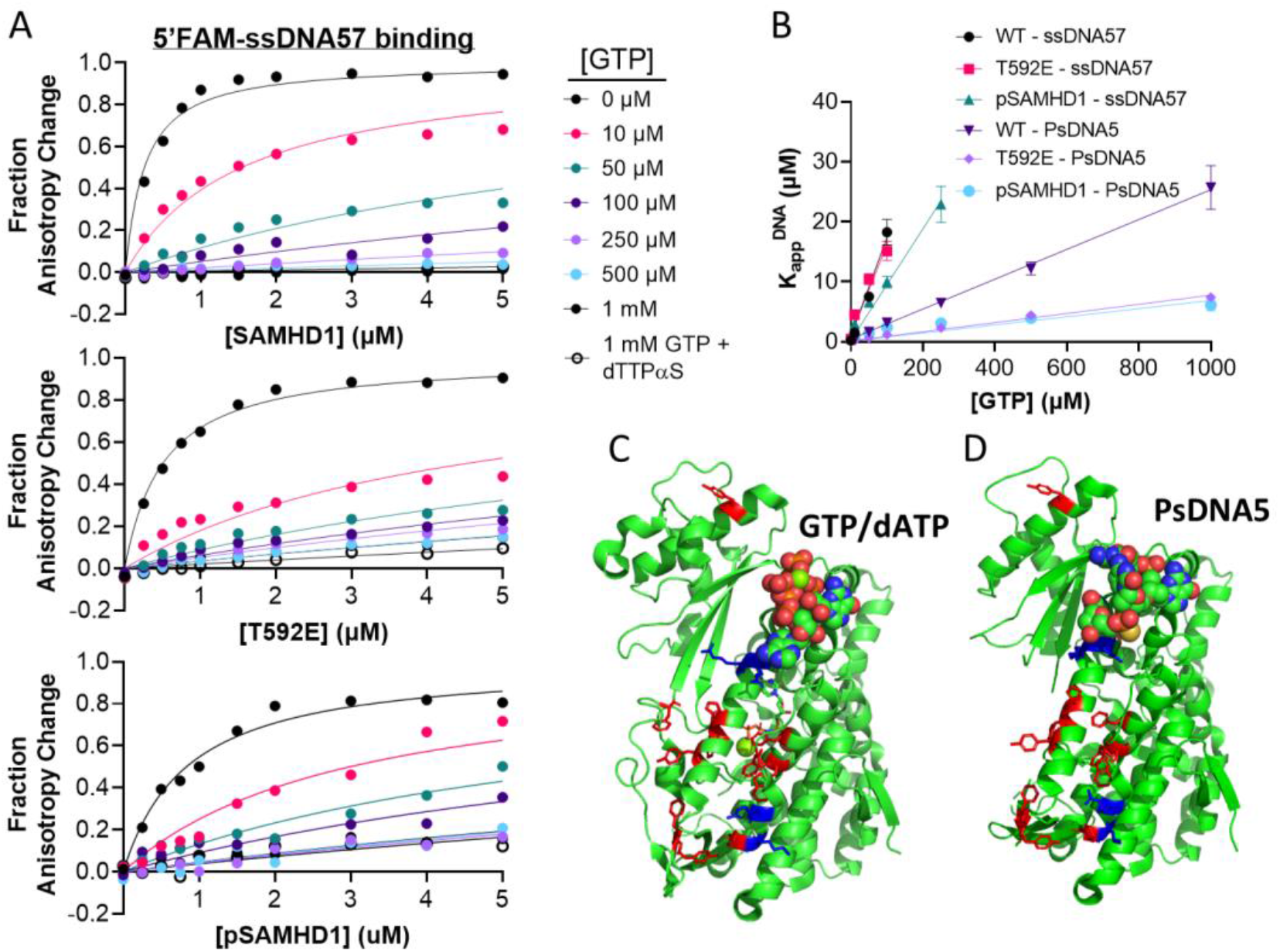
GTP Activator and single-stranded DNA compete for the same binding site on SAMHD1. **(A)** Binding of SAMHD1 wild-type, T592E, and pSAMHD1 to 5’ FAM-labeled ssDNA57 in the presence of no nucleotides, increasing concentrations of GTP, and a combination of GTP/dTTPαS (1 mM each). **(B)** Apparent Kd values from (A) plotted as a function of GTP concentration. Error bars indicate standard error of K_app_^DNA^ from least-squares regression fit of the data in (A) to equation 6. **(C)** Isolated SAMHD1 monomer from the GTP & dATP bound tetrameric SAMHD1 structure (PDB 6TXC). DNA-interacting residues identified via BrdU-crosslinking are highlighted in red and cationic residues involved in DNA binding are highlighted in blue. **(D)** Isolated SAMHD1 monomer from a PsDNA5 bound SAMHD1 structure (PDB 6U6X). DNA-interacting residues identified via BrdU-crosslinking are highlighted in red and cationic residues involved in DNA binding are highlighted in blue.

The similar GTP competition effects observed with PsDNA5 and ssDNA57 suggest that one of the guanines at the 5’-end of ssDNA57 occupies the A1 site. Supporting this binding orientation, our previous BrdU DNA photo-crosslinking approach identified SAMHD1 amino acid residues in the dimer-dimer interface that were near the bound DNA. These residues (red) are mapped on the structure of SAMHD1 in **Figure 2c**, along with cationic residues (blue) that were implicated in ssDNA binding by charge reversal mutations (11). Comparison of the crosslinking results with the structure of PsDNA5 bound to SAMHD1 (**Fig. 2d**) supports the proposal that the 5’ end of longer DNAs would occupy the activator site(s) and extend into the contiguous binding surface identified in the crosslinking experiments. Based on this extended DNA binding site, we surmise that GTP binding to the A1 site increases the mobility of the FAM probe at the 5’-end of ssDNA57, while the remaining DNA chain remains bound to the extended binding site. These data imply that binding of long ssDNA would displace GTP from the A1 site, sterically blocking tetramerization and inhibiting dNTPase activity.

### Effects of T592 variants on binding of GTP and ssDNA to the A1 site

Since GTP is capable of displacing ssDNA, we then asked whether ssDNA could also displace bound GTP. For this question, we validated a new fluorescent probe for the A1 site, N-methyl anthraniloyl (mant) GTP. Mant-GTP is excited at 335 nm and shows an emission maximum at ~440 nm, making it a historically useful environmentally sensitive probe for GTP binding sites (42). We first observed that addition of a near saturating concentration of SAMHD1 (10 μM) to 0.5 μM mant-GTP resulted in an increase in the mant-GTP fluorescence intensity and a 7 nm blueshift in the emission wavelength maximum (**Fig. 3a**), indicating that mant-GTP might be a useful probe of the A1 site. We then established that 1 mM mant-GTP activated the SAMHD1 dTTPase activity in the same manner as an equivalent concentration of GTP (**Fig. 3b, Supplemental Fig. S6**). The specificity of mant-GTP for the A1 site was further established through partial displacement by GTP, but not dATP, which only binds to the A2 and catalytic sites (**Fig. 3c**). Although we observed that addition of 1 mM GTP largely returned the mant-GTP emission maximum to the value observed with free mant-GTP (**Fig. 3a**), consistent with complete competition, the mant-GTP fluorescence intensity increased linearly with SAMHD1 concentration (**Fig. 3c**). This result suggested an additional fluorescence contribution from non-specific binding of mant-GTP. We conclude that the blueshift corresponds to specific binding of mant-GTP to the A1 site (100% competition with GTP), but that the total fluorescence intensity increase had contributions from both specific binding and weak non-specific binding. Accordingly, a standard linear correction for the contribution from non-specific binding was performed on all subsequent measurements using mant-GTP. The ability of mant-GTP to bind to the A1 site and activate the enzyme establishes this ligand as a useful and functional A1 site probe.

**Figure 3.**
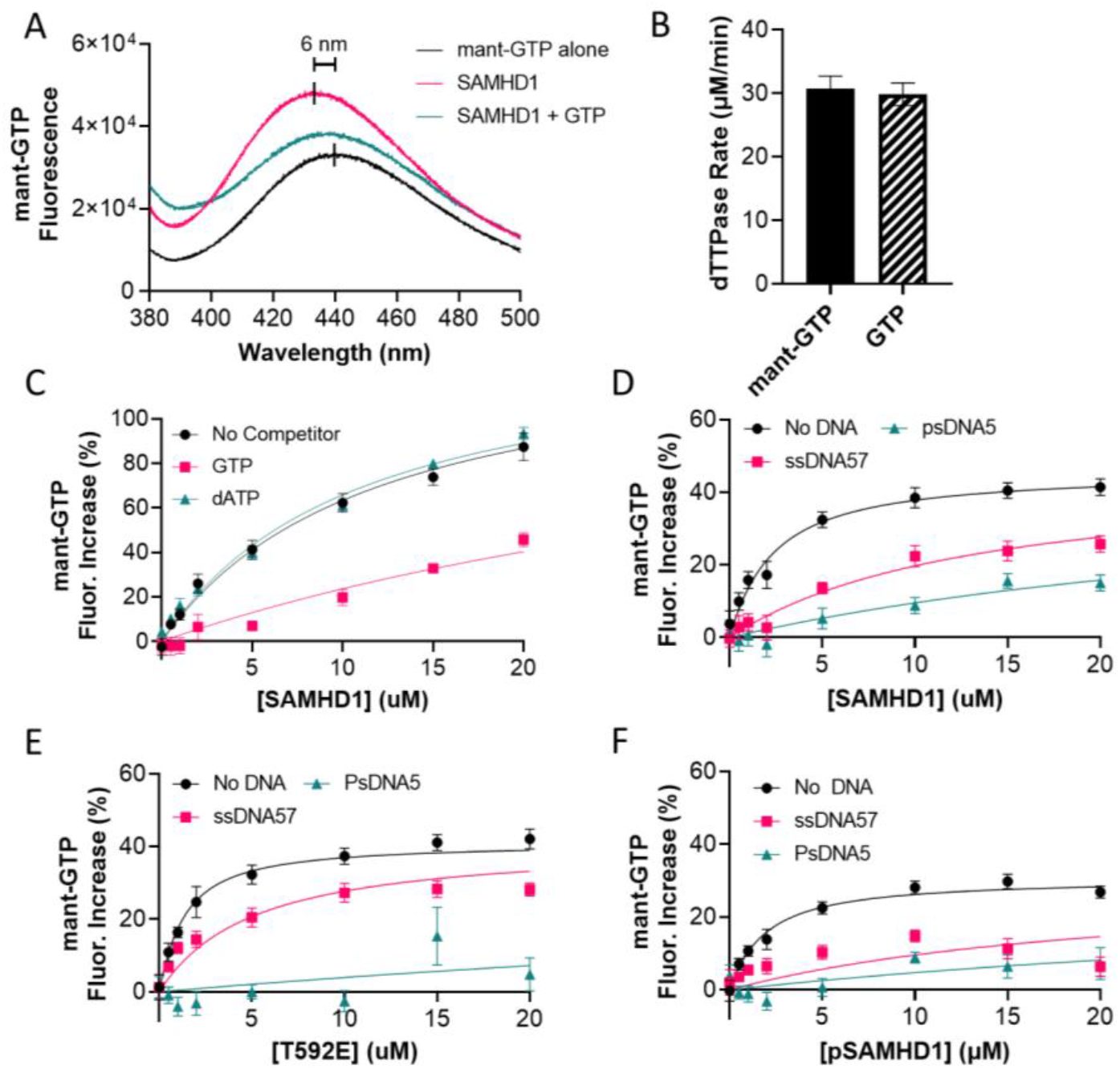
Single-stranded DNA displaces N-methyl anthraniloyl GTP from the A1 site. **(A)** Emission spectra of mant-GTP (0.5 μM) alone and bound to 10 μM SAMHD1. Emission scans were taken from 380 nm to 500 nm with an excitation wavelength of 335 nm. **(B)** Activation of dTTP (1 mM) hydrolysis by mant-GTP (0.5 mM) as compared to the attained under the same concentration of GTP and dTTP from kinetic analysis in fig. 1A. Error bars indicate standard error of reaction rate determined by the linear regression fit of the linear phase of three replicate reactions performed for each condition. (**D)** Binding of wild-type to SAMHD1 mant-GTP (0.5 μM) in the presence of dATP (1 mM) or GTP (1 mM). **(E)** Binding of wild-type SAMHD1 to mant-GTP (0.5 μM) in the presence of ssDNA57 (20 μM) or psDNA5 (20 μM). The nonspecific binding measured in (D) was subtracted from each of the binding isotherms. **(F)** Binding of SAMHD1 T592E to mant-GTP (0.5 μM) in the presence and absence of ssDNA57 (20 μM) or psDNA5 (20 μM). The nonspecific binding measured in Supplemental Fig. S7 was subtracted from each of the binding isotherms. **(G)** Binding of pSAMHD1 to mant-GTP (0.5 μM) in the presence and absence of ssDNA57 (20 μM) or psDNA5 (20 μM). The nonspecific binding baseline attained Fig. S7 was subtracted from each of the binding isotherms. Error bars in (C) through (F) represent standard error of mean from three replicate titrations.

The corrected fluorescence increase associated with the specific binding of mant-GTP to the A1 site was used to obtain dissociation constants of mant-GTP for SAMHD1, T592E, pSAMHD1, (**Fig. 3d, e, f**)(**Supplemental Fig. S7**). The binding affinities and fluorescence amplitude changes were similar for all three of these constructs (*K*_D_ values fell in the range 1 to 2 μM with overlapping confidence intervals). We then asked whether ssDNA57 and PsDNA5 could displace mant-GTP from the A1 site of each of these enzyme forms (**Fig. 3d, e, f**). For each enzyme, PsDNA5 was the most effective competitor, resulting in essentially complete displacement of mant-GTP for T592E and pSAMHD1, and ~65% displacement for wtSAMHD1. Competition was not as efficient for DNA57, even though GTP was found to greatly increase the mobility of the 5’FAM end of DNA57 in the DNA anisotropy measurements (**Fig. 3a**). These differences in apparent competition may arise from conformational differences between bound GTP and mant-GTP. Nevertheless, the two competition experiments with DNA and GTP suggest a linkage between GTP A1 site occupancy and the interaction of the enzyme with the 5’ end of longer DNA molecules.

### Structural basis for communication between T592 phosphorylation and the A1 site

To gain some insights into the structural basis for how ssDNA may gain access to the A1 site, we turned to (i) a coarse-grained normal mode analysis (NMA) of SAMHD1, (ii) a comparison of the structures of the T592E and T592V variants of SAMHD1 via alignment and structure-to-structure morphing using the ChimeraX software (PDB: 4ZWE and 4ZWG, respectively), and (iii) a new cryo-EM structure of SAMHD1 bound to dGTPαS. The NMA was performed on an isolated monomer of SAMHD1 and the relative displacements of individual Cα atoms of the peptide backbone were plotted against the linear sequence for comparison with the observed displacements between the two tetramer crystal structures (**Fig. 4a**). The largest displacements in both analyses were localized in two clustered regions corresponding to residues 462-498 and 555-600. Mapping of Cα NMA fluctuations onto the input SAMHD1 structure revealed that the motion is concentrated in a bundle of α-helixes at the C-terminus of the enzyme which contains T592, hence forth referred to as the C-terminal domain (CtD) (**Fig. 4b**). The excellent correlation between the NMA and the structural changes observed upon phosphomimetic mutation of T592, indicate that perturbations of T592 modulate the intrinsic flexibility of the peptide chain. In other words, the native T592 and its phosphorylated form can stabilize/destabilize different conformations of the CtD that are present in dynamic equilibrium. Videos of the dynamic motions determined from the NMA and the structure morphing approach are found in **Supplemental Videos V1 and V2**.

**Figure 4.**
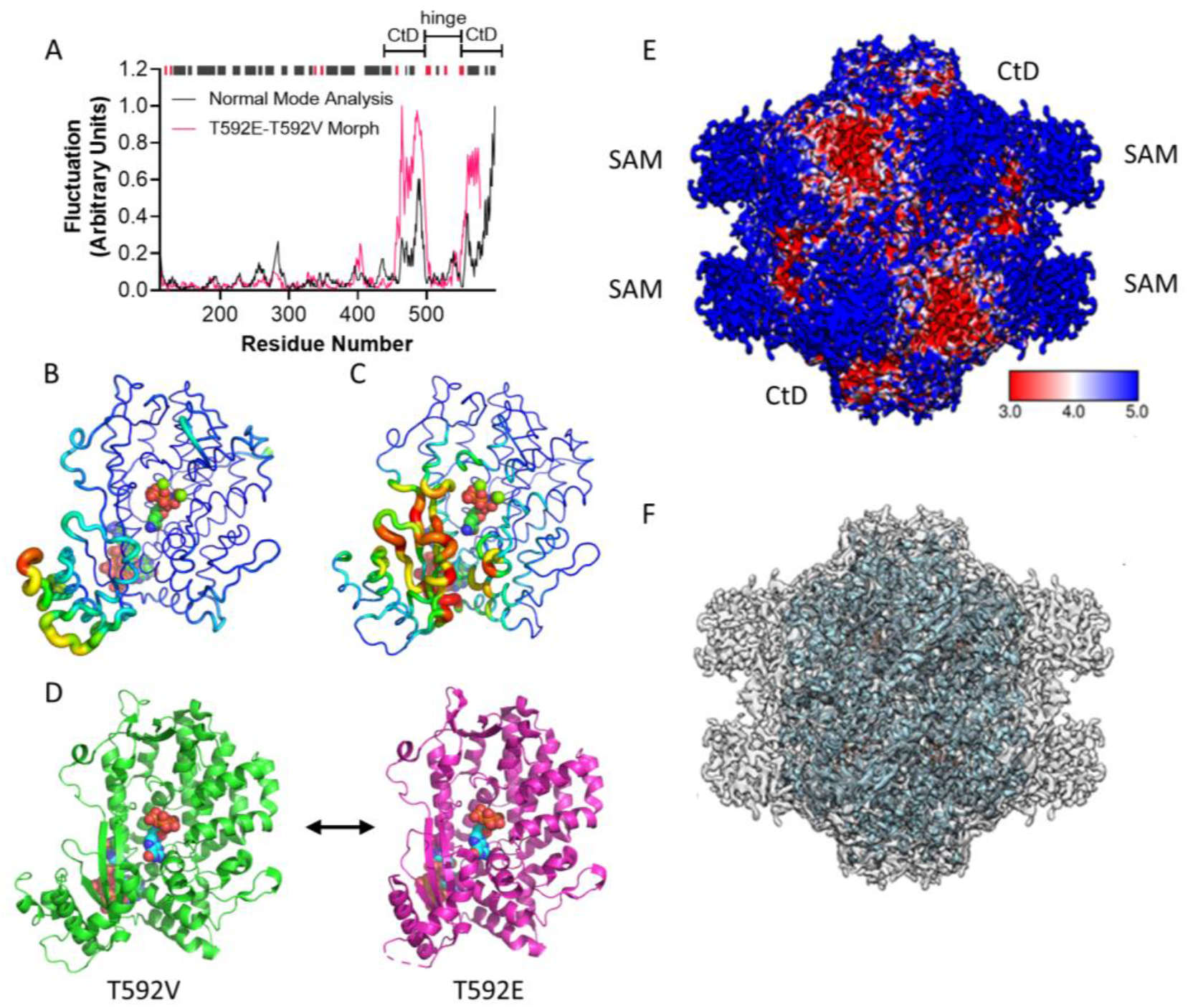
Conformational flexibility of the C-terminal domain. **(A)** Plot of relative fluctuations by residue determined by coarse grained normal mode analysis on an isolated monomer from a tetrameric SAMHD1 structure (PDB: 6TXC) (pink) and by alignment of isolated monomers from tetrameric structures of SAMHD1 T592V (PDB: 4ZWE) and SAMHD1 T592E (PDB: 4ZWG) (black). Secondary structure is indicated at the top of the plot, with black bars indicating α-helices, and red bars indicating β-sheets. The C-terminal Domain (CtD) and flexible hinge region are labeled above the plot. **(B)** Fluctuations determined by normal mode analysis mapped on the structure of SAMHD1 (PDB: 6TXC). **(C)** Deformation energy determined by normal mode analysis mapped on the structure of SAMHD1 (PDB ID 6TXC). **(D)** Conformations of the C-terminal domain (CtD) (T592V – PDB: 4ZWE, T592E – PDB: 4ZWG). The T592E mutation causes the CtD to shift away from the allosteric sites and towards the active site. **(E)** Local resolution cryo-EM map of full-length hSAMHD1 bound to dGTPαS. The cryoEM map is shown at a lower contour to highlight the lower resolution (blue) SAM and C-terminal domains. **(F)** An atomic model based on the SAMHD1 catalytic domain was docked into (E), showing the residual unmodeled densities corresponding to the SAM domains.

A further deconstruction of the NMA and morphing results shows that the CtD lobe can oscillate between two limiting conformations that either (i) partially occlude the catalytic site (stabilized by phosphomimetic mutation T592E), or (ii) partially occlude the A1 site (favored in the T592V structure) (**Fig. 4d**). The motions of the lobe are facilitated by a flexible hinge consisting of two β-strands (498-506, 546-554) flanking the catalytic site (**Fig. 4c**). These findings support a mechanism where intrinsic dynamic motions of the tetramer are perturbed by phosphorylation thereby exposing the A1 site and hindering access to the catalytic site. These effects could easily be attenuated or obscured in steady-state dNTPase kinetic measurements where the rate-limiting steps are much slower than these events, as well as in equilibrium binding measurements that are heavily weighted towards detection of the major bound species and not changes in populations of minor species that are in dynamic equilibrium.

To bolster these findings, we solved a structure of tetrameric SAMHD1 bound to dGTPαS via cryo-EM. Examination of the density map colored via local resolution shows that tetrameric SAMHD1 consists of a high-resolution core with eight low-resolution (i.e., conformationally flexible) lobes on the surface of the tetramer (**Fig. 4e**) (**Supplemental Fig. S8**). An atomic model confidently built into the SAMHD1 catalytic domain of the cryo-EM map using published crystal structures (PDB: 4TNX) as a guide (**Fig. 4f**) allows definitive assignment of four lobes as the CtD, with the remaining four inferred to be the SAM domains. Examination of the structure indicates that CtD and SAM domains of adjacent chains are spatially close together in the context of the tetrameric holoenzyme, with pairs of allosteric sites buried in the cleft underneath the two. This highlights potentially important functional roles for the CtD and SAM domains in maintaining the stability of the tetramer and controlling access to the allosteric sites and nucleic acid binding site at the tetramer interface.

### Tetrameric state of T592 variants has a reduced thermal melt temperature

Thermal shift assays utilizing fluorogenic dyes that bind to exposed hydrophobic surfaces of proteins are a convenient and efficient tool for evaluating changes in protein thermal stability arising from mutation or ligand binding (43). We used thermal shift assays to evaluate the effects of phosphomimetic mutation, phosphorylation and two C-terminal truncations (Δ583-626 and Δ600-626) on the thermal stability of SAMHD1 tetramer using the common fluorescent dye SYPRO orange. The deletion variants differ in that Δ583-626 lacks T592 while Δ600-626 retains this residue. We first determined the melting temperature (*T*_m_) of each enzyme in the absence of nucleotide and found that the *T_m_* of all enzymes fell between 44 and 46 °C with no clear correlation between *T_m_* and perturbation of T592 (**Supplemental Fig. S9**). The tetrameric form of each enzyme was generated by the addition of a saturating concentration of dGTPαS (2 mM), which binds to the A1, A2 and the catalytic sites (4, 44). In comparison to the unliganded enzymes, the tetramer forms all showed a remarkable increase in *T_m_*, now falling in a range of 74 to 78 °C (1^st^ derivative plots are shown in **Fig. 5a**). Except for variant Δ600-626, each showed a reduced *T*_m_ value compared to SAMHD1, indicating that the tetramers were destabilized (**Fig. 5b**). The *T*_m_ value of SAMHD1 tetramer in the presence of dGTPαS was 77.07 ± 0.11 °C, while the phosphomimetic T592E and pSAMHD1 had *T*_m_ values that were both 1.2 °C lower (*T_m_^T592E^* = 75.98 ± 0.16 °C, *T_m_^pSAMHD1^* = 76.01 ± 0.25 °C). The *T*_m_ value for Δ583-626 was nearly 3 degrees lower than full-length SAMHD1 (*T*_m_ = 74.30 ± 0.19 °C), while Δ600-626 was slightly elevated (*T*_m_ = 77.65 ± 0.06 °C). These results indicate that T592 phosphorylation destabilizes the nucleotide bound tetrameric state of SAMHD1, while having little impact on the monomeric apoenzyme. The mechanism of destabilization appears involve a stabilizing interaction of the T592 hydroxyl side chain that is disrupted upon phosphorylation or mutation. Supporting this view, the hydroxyl group of Thr-592 participates in a hydrogen bonding triad with Asp-585 and Lys-580 (**Fig. 5c**), which would be disrupted upon phosphorylation of T592 based on charge repulsion and steric clashes. The alternative explanation where phosphomimetic mutation or phosphorylation destabilizes the tetramer is not consistent with the lower thermal melting temperature of Δ583-626 which lacks T592.

**Figure 5.**
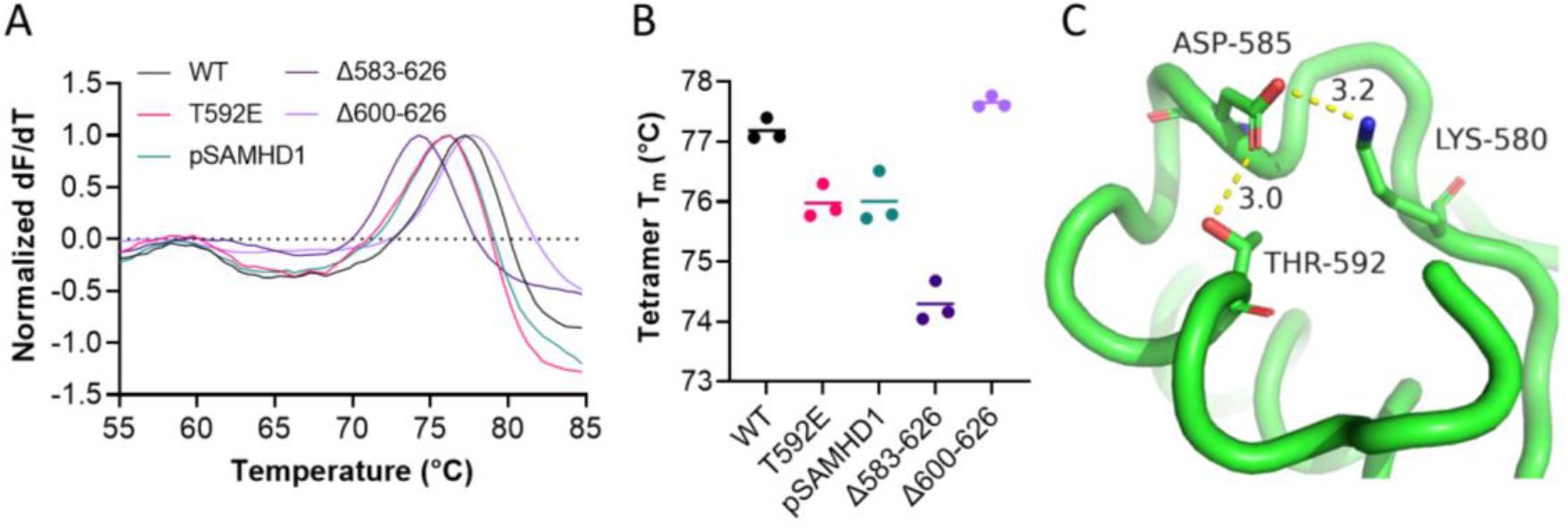
Phosphorylation, phosphomimetic mutation, and elimination of the phosphorylation site decrease the thermal stability the SAMHD1 tetramer. **(A)** Normalized first derivative plot of thermal melts of wild-type SAMHD1, T592E, pSAMHD1, SAMHD1 Δ583-626, and SAMHD1 Δ600-626. SAMHD1 (3 μM) was incubated with 2 mM dGTPαS to drive the system to a tetrameric state. Temperature was ramped up from 25 °C to 85 °C. **(B)** Melt temperatures of individual replicates from (A) calculated in TSA-craft. **(C)** Hydrogen bonding network formed by threonine 592, aspartate 585, and lysine 580 (PDB: 6TXC).

### Tetrameric state of the T592 variants is susceptible to disruption by ssDNA

We previously developed a method to “pre-activate” SAMHD1 by incubating it in the presence of high concentration of GTP activator and then diluting the activated enzyme into a reaction buffer containing a substrate dNTP but no activator (4). This approach (variations of which have been subsequently used by other groups), allows the activation step to be separated from enzyme turnover because activating nucleotides remain tightly bound after dilution of the tetramer into reaction buffer containing substrate. Under such conditions in the presence of substrate, the enzyme tetramer persists for at least 6 h with no change in activity.

Here we explored the behaviors of pre-activated SAMHD1, T592E and pSAMHD1 when each were diluted one-hundred-fold into a reaction buffer *without substrate* in the presence and absence of 50 nM 5’FAM-ssDNA57 to monitor DNA binding through the increase in anisotropy (**Fig. 6a**). This experiment is designed to explore if the dynamic behavior of the T592E and pSAMHD1 enzymes facilitates invasion of ssDNA into the dimer interface of the tetramer. For SAMHD1, T592E and pSAMHD1, simply diluting the enzymes into buffer containing 5’FAM-ssDNA57 results in immediate attainment of the higher anisotropy value expected from binding 50 nM DNA (**Fig. 2b, c, d**) (dead time ~ 15s). In contrast, when the enzymes are pre-activated in the presence of 1 mM dGTP and then diluted one hundred-fold into a buffer containing 50 nM 5’FAM-ssDNA57, a slower increase in anisotropy occurs that is consistent with the requirement to disassemble the tetramer prior to or during binding of the DNA. The rates of DNA binding followed the trend SAMHD1 (*t*_1/2_ = 134 s) < T592E (*t*_1/2_ = 36 s) < pSAMHD1 (*t*_1/2_ = 16 s). Thus, pSAMHD1 binds DNA about 8-fold faster as compared to the non-phosphorylated enzyme. Each of the kinetic curves was repeated five times in independent measurements and the average values and standard errors are shown in Figure 5.

**Figure 6.**
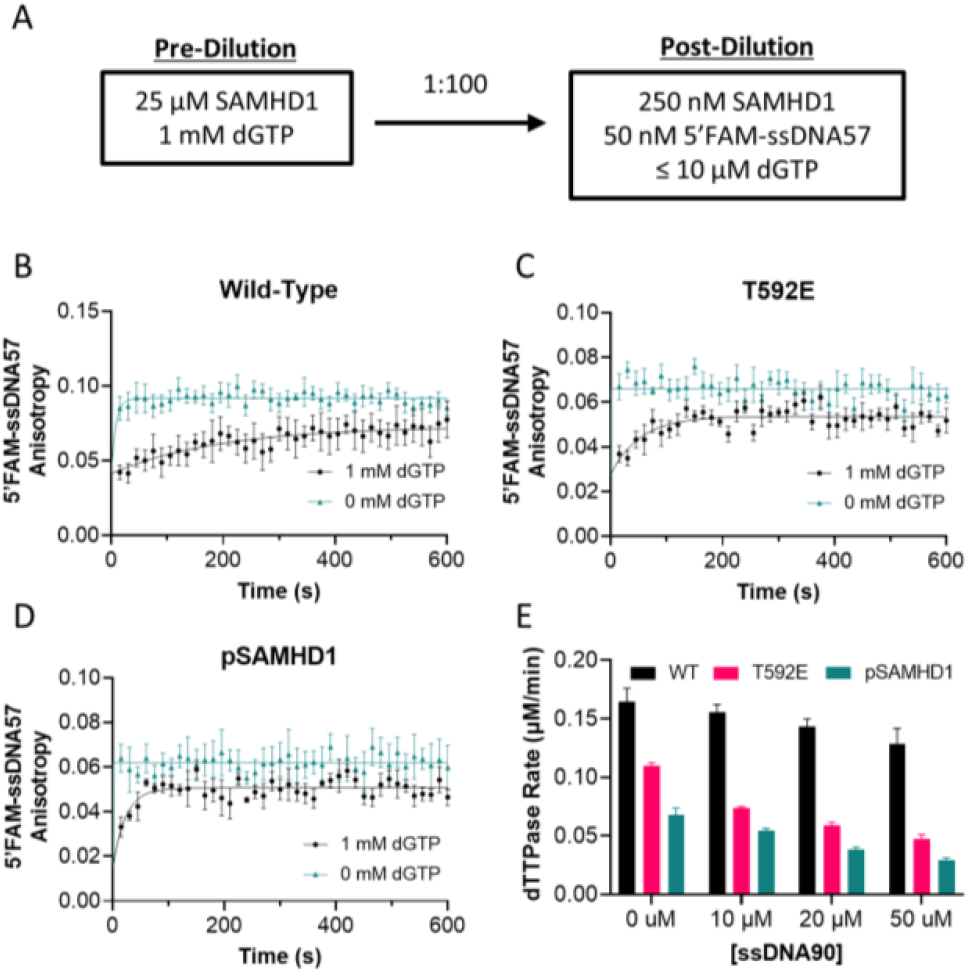
Both phosphorylation and phosphomimetic mutation sensitize SAMHD1 to inhibition by single-stranded DNA. **(A)** Schematic of dilution experiment in (B), (C), and (D). A 25 μM SAMHD1 solution was preincubated for 30 seconds with 1 mM dGTP or with no dGTP, then diluted 100-fold into a solution of 50 nM 5’FAM labeled ssDNA57. Anisotropy of the FAM fluorophore was measured at 15 second intervals for 10 minutes. **(B)** ssDNA binding kinetics of SAMHD1 WT. **(C)** ssDNA binding kinetics of SAMHD1 T592E. **(D)** ssDNA binding kinetics of pSAMHD1. Error bars for (B), (C), and (D) indicate standard error of mean from 5 replicate reactions. **(E)** Inhibition of SAMHD1 dNTPase activity by increasing concentrations of single-stranded DNA. SAMHD1 (0.5 μM) was incubated with 10 μM dTTP, 10 μM GTP, and 0-50 μM ssDNA90. Error bars represent standard error of the reaction rate as determined by linear regression analysis of the linear phase of three replicate reactions.

Since binding of long ssDNA requires access to the dimer interface, the above results suggested that the dNTPase activity of T592E and pSAMHD1 would also be inhibited in the presence of ssDNA under similar conditions of limiting nucleotides. To test this prediction, we carried out dNTPase assays for the three enzymes (0.5 μM) using 10 μM dTTP, 10 μM GTP and increasing concentrations of ssDNA90 in the range 0 to 50 μM (**Fig. 6e**). The dTTP substrate concentration used in this experiment is typical of that in many human cell lines (45). Consistent with the prediction, the dNTPase rates of T592E and pSAMHD1 declined in response to increasing DNA concentrations while SAMHD1 remained largely unaffected by the presence of DNA. Taken together, these findings suggest that phosphorylation destabilizes the tetramer, promotes ssDNA binding land leads to selective inhibition of pSAMHD1 as compared to SAMHD1.

### Tetramer dissociation kinetics of SAMHD1 and its T592 variants

To directly establish that the presence of T592 in wild-type SAMHD1 promotes stabilization of the tetramer and resistance to ssDNA invasion, we monitored tetramer dissociation in the absence and presence of ssDNA using a previously described glutaraldehyde crosslinking assay (GAXL) that allows interrogation of the oligomeric states of SAMHD1 that are present under various solution conditions (4, 46). The experimental approach mimics the pre-activation and dilution experiments described above (**Fig. 7a**). In the first series of experiments, we mixed 25 μM of SAMHD1 or the variants T592E, pSAMHD1, Δ583-626 and Δ600-626 with 1 mM dGTP before rapidly diluting one hundred-fold into reaction buffer (final [enzyme] = 250 nM). As a function of time post-dilution, samples of each reaction were removed and rapidly crosslinked using 50 mM glutaraldehyde. The fraction tetramers present at each time were quantified by fluorescence imaging after separation by SDS-PAGE and staining using Coomassie dye. In the absence of ssDNA, the variants where T592 was deleted, mutated to glutamate or phosphorylated showed dramatically increased rates of tetramer (T) dissociation into monomers (M) as compared to native SAMHD1 and Δ600-626 which both retain T592 (**Fig. 7b**). Although the tetramer dissociations were best fit to double exponential decays, a simple comparison of the phenomenological half-life for dissociation of SAMHD1 (~55.6 min), indicates that dissociation occurs at least 6 to 15-fold slower than T592E, pSAMHD1, and Δ583-626 (~ 8.8, 3.6 and 4.2 min, respectively). Like the thermal shift results, these findings indicate that T592 is involved in tetramer stabilization which is abrogated by phosphomimetic mutation, phosphorylation, and by elimination of the phosphorylation site. Interestingly, Δ600-626 appears to form a moderately more stable tetramer than the wild-type, with a phenomenological half-life of 260 min, consistent with its higher melting temperature in the TSA measurements.

**Figure 7.**
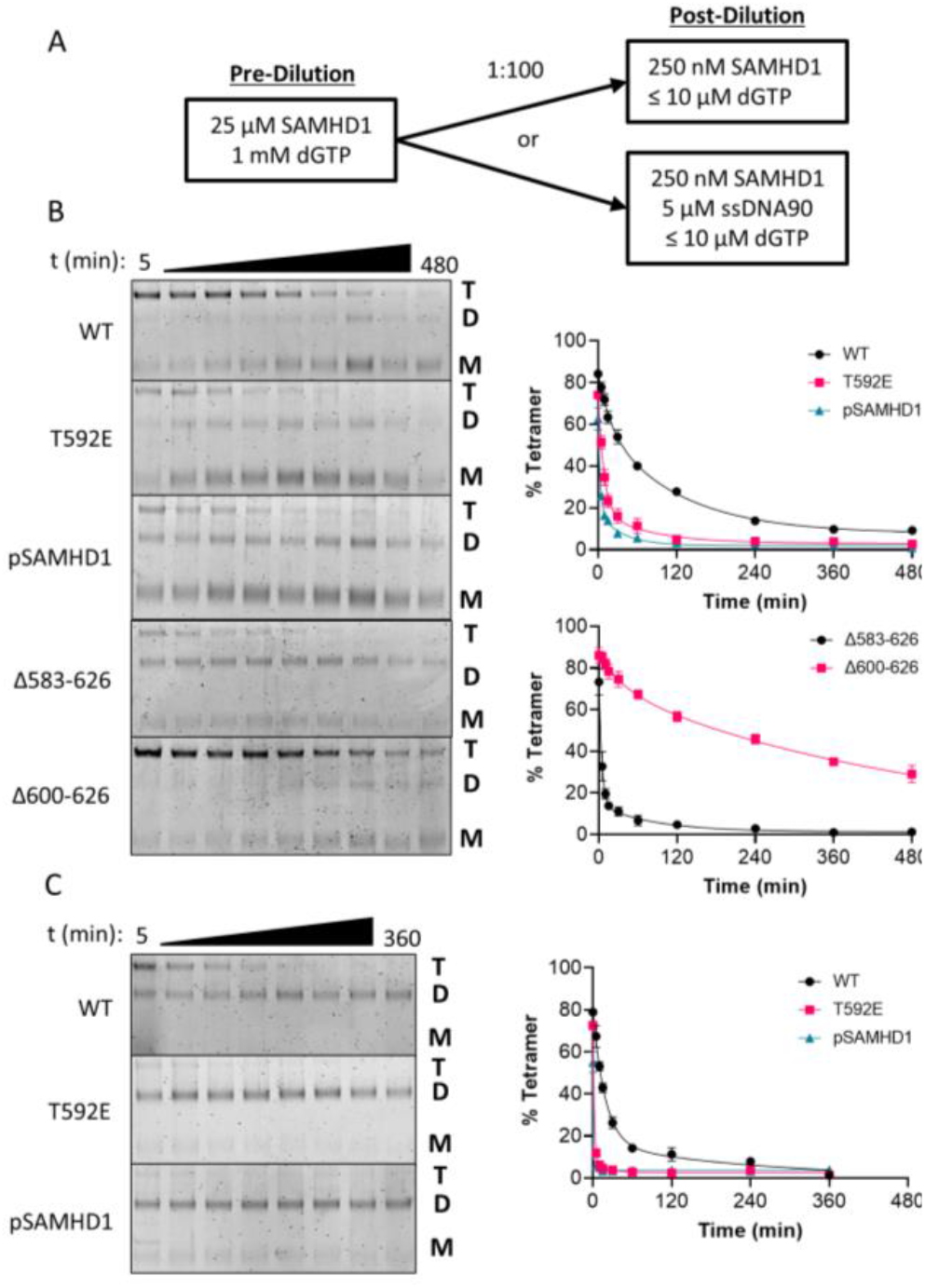
SAMHD1 tetramer dissociation is enhanced by phosphomimetic mutation, phosphorylation, elimination of the phosphorylation site, and by the presence of single-stranded DNA. **(A)** Schematic of dilution experiment. SAMHD1 (25 μM) was incubated with 1 mM dGTP for 30 seconds, then rapidly diluted 100-fold into a solution containing no activating nucleotides, as in (B), or into a solution of 5 μM ssDNA90, as in (C), and crosslinked in glutaraldehyde (50 mM) at regular timer intervals to monitor dissociation of the tetrameric state. **(B)** Dissociation kinetics of wild-type SAMHD1, T592E, pSAMHD1, SAMHD1 Δ583-626, and SAMHD1 Δ600-626. Select excerpts of gels are shown on the left, and quantification of the three replicates done for each enzyme are shown on the right. Error bars represent standard error of mean from three replicate reactions. **(C)** Dissociation kinetics of wild-type SAMHD1, T592E, pSAMHD1 in the presence of 5 μM ssDNA90. Select excerpts of gels are shown on the left, and quantification of the three replicates done for each enzyme are shown on the right. Error bars indicate standard error of mean for three replicate reactions.

The same tetramer dissociation experiment was repeated except that the dilution buffer contained 5 μM ssDNA57 (**Fig. 7c**). The presence of ssDNA was observed to accelerate tetramer dissociation for all enzymes, but once again, the most rapid dissociation was observed with the T592 variants. Dissociation of SAMHD1 and T592E tetramers was accelerated ~3 and 5-fold in the presence of ssDNA (*t*_1/2_ = 17 and 1.6 min, respectively), while 90% of pSAMHD1 tetramer dissociated in the first five minutes of the experiment, when the first time point was taken. Of interest, the final crosslinked species in the presence of ssDNA migrated predominantly as dimers, whereas in the absence of DNA, a mixture of monomers and dimers were produced. These findings indicate that ssDNA lowers the activation barrier for the rate-limiting step in tetramer dissociation and that DNA binding brings two monomers in proximity for crosslinking. The lack of glutaraldehyde crosslinking between the enzyme and DNA is not unexpected because the exo-cyclic amine groups in DNA bases are far less reactive than primary amines in lysine side chains of proteins. The binding of two SAMHD1 monomers to DNA of this length is consistent with previous AFM images and binding stoichiometry measurements (12).

## DISCUSSION

Although functional requirements for SAMHD1 T592 phosphorylation in restart of stalled replication forks and promoting DNA end resection in homologous recombination have been reported (14, 21), there is no coherent mechanism for how this covalent modification changes SAMHD1 activity and facilitates such DNA repair processes. Consistent with previous reports, we have shown that phosphorylation has unremarkable effects on dNTPase activity. Since the DNA binding surface of SAMHD1 is not exposed in its tetrameric form, suggests that a productive role for phosphorylation during ssDNA transactions would be to increase exposure of the DNA binding surface (12). Our findings suggest a bi-partite role for phosphorylation in inactivating SAMHD1 dNTPase and promoting its association with ssDNA during these repair transactions: tetramer destabilization by phosphorylation, followed by ssDNA trapping of the A1 site and invasion into the dimer-dimer DNA binding interface of the tetramer. This mechanism provides regiospecificity because it requires both phosphorylation and free ssDNA.

The extensive series of in vitro experiments performed in this study have established the required elements for such a mechanism (**Fig. 8**). A central paradox is how a modification that gives rise to unremarkable changes in standard activity measurements of SAMHD1 can serve as a functional switch to turn on the S phase replication fork activity of SAMHD1. Our thermal shift results, ssDNA trapping, structural analyses, and observations that ssDNA and GTP bind antagonistically to the A1 site all indicate that removal or phospho-modification of Thr592 energetically destabilizes the SAMHD1 tetramer leading to ssDNA invasion of the tetramer interface. This mechanism involves several steps that are supported by the collective data (**Figure 8**). The thermal melt shifts of tetrameric SAMHD1 upon phosphorylation indicate that this modification increases the free energy of the tetramer (destabilizes) through negation of the favorable interactions of the Thr592 side chain (step 1). This destabilization shifts the internal conformational equilibrium of the CtD such that the A1 site is more exposed, and the catalytic site is more occluded (**Fig. 4**). This conformational state is minimally populated in the unphosphorylated enzyme due to the stabilization provided by Thr592. However, in the phosphorylated enzyme the internal equilibrium shifts to increase the concentration of the unstable conformation, and the more exposed A1 site can be accessed by guanine nucleotides present within ssDNA (step 2). The nascent interaction with ssDNA at the A1 site then promotes further invasion of ssDNA into its previously defined extended binding site along the tetramer interface. Since the presence of ssDNA accelerates tetramer dissociation (**Fig. 7**), the activation barrier is decreased in the presence of ssDNA (step 2). The crosslinking results suggest that with ssDNA90, dimers of SAMHD1 are bound to DNA and are trapped by chemical crosslinking. Contrary to previous reports, we see no evidence of mixed-occupancy tetramer formation in the presence of ssDNA (41). In the absence of ssDNA, tetramer dissociation is much slower and results in a mixture of dimers and monomers (**Fig. 7**). A key feature of the above mechanism is the invasion of ssDNA using a guanine specific DNA interaction with the A1 site. This is supported by the crystal structure and the displacement of GTP by both PsDNA5 and ssDNA57. Such a mechanism would provide specificity for ssDNA because guanine bases are not accessible in duplex DNA. Indeed, SAMHD1 has poor affinity for duplex DNA. A full investigation into DNA length and sequence effects on this invasion mechanism will be reported in a subsequent study.

**Figure 8.**
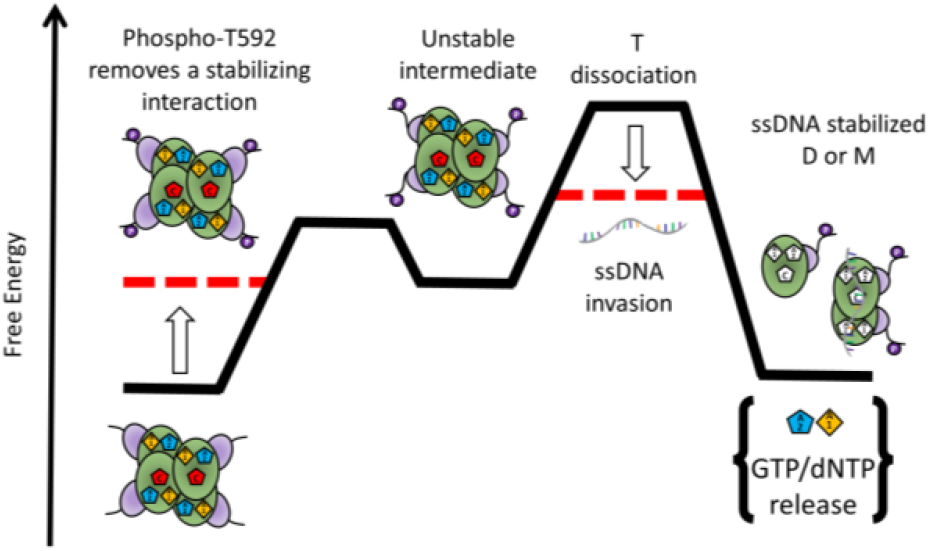
A model for phosphorylation-dependent disassembly of SAMHD1 at stalled replication forks. Structural perturbation of the C-terminal domain (shown in purple) caused by phosphorylation decreases the stability of the tetrameric state. While this destabilization does not have a strong impact on the steady-state dNTPase activity of the enzyme in the presence of activating and substrate nucleotides, it does increase tetramer dynamics. The presence of ssDNA induces rapid dissociation of the pSAMHD1 tetramer into predominantly dimeric products. Due to the antagonistic nature of GTP and ssDNA binding, we presume that dissociation involves displacement of GTP by ssDNA strand invasion into the allosteric sites and DNA binding interface.

Our in vitro studies show how the dynamic properties of pSAMHD1 are more suited for rapid displacement of GTP followed by ssDNA strand invasion than native SAMHD1. If these properties are manifested at replication forks and sites of homologous recombination, they suggest that SAMHD1 tetramer is disassembled during such transactions. Although we focus on the aspect of ssDNA binding, these DNA interactions do not negate the possibility of SAMHD1 binding to other proteins at these sites such as C-terminal interacting protein (CTIP) and MRE-11 as previously reported (14, 21).

## DATA AVAILABILITY

All data pertaining to this study are available by request to the corresponding author. The final SAMHD1-dGTPαS complex cryo-EM density map and model are deposited in the Electron Microscopy Data Bank (EMDB) under accession code EMD-26567 and Protein Data Bank (PDB) under accession code 7UJN, respectively.

## Author contributions

BO: conceived study, designed, executed all kinetic, thermodynamic, and biochemical experiments, and wrote the paper

KWH: cryo-EM data processing, model building and structural analysis

MA: grid preparation, sample purification and characterization, grid screening, and data collection

SH: structural analysis

BBol: MS

JC: SAMHD1 phosphorylation

MP: SAMHD1 phosphorylation

BBos and DS: obtained funding, manuscript proofing

JTS: obtained funding, conceived study, wrote the paper

## ACKNOWLEDGEMENTS

We thank You-Ai He for providing purified CDK2/cyclin E1, as well as Todd VanArsdale and James E. Solowiej for initial thoughts and studies on SAMHD1 phosphorylation.

## References

1. Knecht, K.M., Buzovetsky, O., Schneider, C., Thomas, D., Srikanth, V., Kaderali, L., Tofoleanu, F., Reiss, K., Ferreirós, N., Geisslinger, G., et al. (2018) The structural basis for cancer drug interactions with the catalytic and allosteric sites of SAMHD1. Proc. Natl. Acad. Sci. U. S. A., 115, E10022–E10031.

2. Jang, S., Zhou, X. and Ahn, J. (2016) Substrate Specificity of SAMHD1 Triphosphohydrolase Activity Is Controlled by Deoxyribonucleoside Triphosphates and Phosphorylation at Thr592. Biochemistry, 55, 5635–5646.

3. Morris, E.R. and Taylor, I.A. (2019) The missing link: allostery and catalysis in the anti-viral protein SAMHD1. Biochem. Soc. Trans., 47, 1013–1027.

4. Hansen, E.C., Seamon, K.J., Cravens, S.L. and Stivers, J.T. (2014) GTP activator and dNTP substrates of HIV-1 restriction factor SAMHD1 generate a long-lived activated state. Proc. Natl. Acad. Sci. U. S. A., 111, E1843–51.

5. Ji, X., Tang, C., Zhao, Q., Wang, W. and Xiong, Y. (2014) Structural basis of cellular dNTP regulation by SAMHD1. Proc. Natl. Acad. Sci. U. S. A., 111, E4305–14.

6. Schwefel, D., Groom, H.C.T., Boucherit, V.C., Christodoulou, E., Walker, P.A., Stoye, J.P., Bishop, K.N. and Taylor, I.A. (2014) Structural basis of lentiviral subversion of a cellular protein degradation pathway. Nature, 505, 234–238.

7. Chen, S., Bonifati, S., Qin, Z., St Gelais, C., Kodigepalli, K.M., Barrett, B.S., Kim, S.H., Antonucci, J.M., Ladner, K.J., Buzovetsky, O., et al. (2018) SAMHD1 suppresses innate immune responses to viral infections and inflammatory stimuli by inhibiting the NF-κB and interferon pathways. Proc. Natl. Acad. Sci. U. S. A., 115, E3798–E3807.

8. Hrecka, K., Hao, C., Gierszewska, M., Swanson, S.K., Kesik-Brodacka, M., Srivastava, S., Florens, L., Washburn, M.P. and Skowronski, J. (2011) Vpx relieves inhibition of HIV-1 infection of macrophages mediated by the SAMHD1 protein. Nature, 474, 658–661.

9. Schwefel, D., Boucherit, V.C., Christodoulou, E., Walker, P.A., Stoye, J.P., Bishop, K.N. and Taylor, I.A. (2015) Molecular determinants for recognition of divergent SAMHD1 proteins by the lentiviral accessory protein Vpx. Cell Host Microbe, 17, 489–499.

10. Laguette, N., Rahm, N., Sobhian, B., Chable-Bessia, C., Münch, J., Snoeck, J., Sauter, D., Switzer, W.M., Heneine, W., Kirchhoff, F., et al. (2012) Evolutionary and functional analyses of the interaction between the myeloid restriction factor SAMHD1 and the lentiviral Vpx protein. Cell Host Microbe, 11, 205–217.

11. Seamon, K.J., Bumpus, N.N. and Stivers, J.T. (2016) Single-Stranded Nucleic Acids Bind to the Tetramer Interface of SAMHD1 and Prevent Formation of the Catalytic Homotetramer. Biochemistry, 55, 6087–6099.

12. Seamon, K.J., Sun, Z., Shlyakhtenko, L.S., Lyubchenko, Y.L. and Stivers, J.T. (2015) SAMHD1 is a single-stranded nucleic acid binding protein with no active site-associated nuclease activity. Nucleic Acids Res., 43, 6486–6499.

13. Coquel, F., Neumayer, C., Lin, Y.-L. and Pasero, P. (2018) SAMHD1 and the innate immune response to cytosolic DNA during DNA replication. Curr. Opin. Immunol., 56, 24–30.

14. Daddacha, W., Koyen, A.E., Bastien, A.J., Head, P.E., Dhere, V.R., Nabeta, G.N., Connolly, E.C., Werner, E., Madden, M.Z., Daly, M.B., et al. (2017) SAMHD1 Promotes DNA End Resection to Facilitate DNA Repair by Homologous Recombination. Cell Rep., 20, 1921–1935.

15. Cabello-Lobato, M.J., Wang, S. and Schmidt, C.K. (2017) SAMHD1 Sheds Moonlight on DNA Double-Strand Break Repair. Trends Genet., 33, 895–897.

16. Batalis, S., Rogers, L.C., Hemphill, W.O., Mauney, C.H., Ornelles, D.A. and Hollis, T. (2021) SAMHD1 Phosphorylation at T592 Regulates Cellular Localization and S-phase Progression. Front Mol Biosci, 8, 724870.

17. Arnold, L.H., Groom, H.C.T., Kunzelmann, S., Schwefel, D., Caswell, S.J., Ordonez, P., Mann, M.C., Rueschenbaum, S., Goldstone, D.C., Pennell, S., et al. (2015) Phospho-dependent Regulation of SAMHD1 Oligomerisation Couples Catalysis and Restriction. PLoS Pathog., 11, e1005194.

18. Bhattacharya, A., Wang, Z., White, T., Buffone, C., Nguyen, L.A., Shepard, C.N., Kim, B., Demeler, B., Diaz-Griffero, F. and Ivanov, D.N. (2016) Effects of T592 phosphomimetic mutations on tetramer stability and dNTPase activity of SAMHD1 can not explain the retroviral restriction defect. Sci. Rep., 6, 31353.

19. White, T.E., Brandariz-Nuñez, A., Valle-Casuso, J.C., Amie, S., Nguyen, L.A., Kim, B., Tuzova, M. and Diaz-Griffero, F. (2013) The retroviral restriction ability of SAMHD1, but not its deoxynucleotide triphosphohydrolase activity, is regulated by phosphorylation. Cell Host Microbe, 13, 441–451.

20. Tramentozzi, E., Ferraro, P., Hossain, M., Stillman, B., Bianchi, V. and Pontarin, G. (2018) The dNTP triphosphohydrolase activity of SAMHD1 persists during S-phase when the enzyme is phosphorylated at T592. Cell Cycle, 17, 1102–1114.

21. Coquel, F., Silva, M.-J., Técher, H., Zadorozhny, K., Sharma, S., Nieminuszczy, J., Mettling, C., Dardillac, E., Barthe, A., Schmitz, A.-L., et al. (2018) SAMHD1 acts at stalled replication forks to prevent interferon induction. Nature, 557, 57–61.

22. Maelfait, J., Bridgeman, A., Benlahrech, A., Cursi, C. and Rehwinkel, J. (2016) Restriction by SAMHD1 Limits cGAS/STING-Dependent Innate and Adaptive Immune Responses to HIV-1. Cell Rep., 16, 1492–1501.

23. Mlcochova, P., Sutherland, K.A., Watters, S.A., Bertoli, C., de Bruin, R.A., Rehwinkel, J., Neil, S.J., Lenzi, G.M., Kim, B., Khwaja, A., et al. (2017) A G1-like state allows HIV-1 to bypass SAMHD1 restriction in macrophages. EMBO J., 36, 604–616.

24. Mlcochova, P., Winstone, H., Zuliani-Alvarez, L. and Gupta, R.K. (2020) TLR4-Mediated Pathway Triggers Interferon-Independent G0 Arrest and Antiviral SAMHD1 Activity in Macrophages. Cell Rep., 30, 3972–3980.e5.

25. Ferreira, I.A.T.M., Porterfield, J.Z., Gupta, R.K. and Mlcochova, P. (2020) Cell Cycle Regulation in Macrophages and Susceptibility to HIV-1. Viruses, 12, 839.

26. Yan, J., Hao, C., DeLucia, M., Swanson, S., Florens, L., Washburn, M.P., Ahn, J. and Skowronski, J. (2015) CyclinA2-Cyclin-dependent Kinase Regulates SAMHD1 Protein Phosphohydrolase Domain. J. Biol. Chem., 290, 13279–13292.

27. Tang, C., Ji, X., Wu, L. and Xiong, Y. (2015) Impaired dNTPase activity of SAMHD1 by phosphomimetic mutation of Thr-592. J. Biol. Chem., 290, 26352–26359.

28. Wang, Z., Bhattacharya, A., Villacorta, J., Diaz-Griffero, F. and Ivanov, D.N. (2016) Allosteric Activation of SAMHD1 Protein by Deoxynucleotide Triphosphate (dNTP)-dependent Tetramerization Requires dNTP Concentrations That Are Similar to dNTP Concentrations Observed in Cycling T Cells. J. Biol. Chem., 291, 21407–21413.

29. Lee, P.-H., Huang, X.X., Teh, B.T. and Ng, L.-M. (2019) TSA-CRAFT: A Free Software for Automatic and Robust Thermal Shift Assay Data Analysis. SLAS Discov, 24, 606–612.

30. Morris, E.R., Caswell, S.J., Kunzelmann, S., Arnold, L.H., Purkiss, A.G., Kelly, G. and Taylor, I.A. (2020) Crystal structures of SAMHD1 inhibitor complexes reveal the mechanism of water-mediated dNTP hydrolysis. Nat. Commun., 11, 3165.

31. Grant, B.J., Rodrigues, A.P.C., ElSawy, K.M., McCammon, J.A. and Caves, L.S.D. (2006) Bio3d: an R package for the comparative analysis of protein structures. Bioinformatics, 22, 2695–2696.

32. Pettersen, E.F., Goddard, T.D., Huang, C.C., Meng, E.C., Couch, G.S., Croll, T.I., Morris, J.H. and Ferrin, T.E. (2021) UCSF ChimeraX: Structure visualization for researchers, educators, and developers. Protein Sci., 30, 70–82.

33. Zivanov, J., Nakane, T., Forsberg, B.O., Kimanius, D., Hagen, W.J., Lindahl, E. and Scheres, S.H. (2018) New tools for automated high-resolution cryo-EM structure determination in RELION-3. Elife, 7.

34. Mastronarde, D.N. (2005) Automated electron microscope tomography using robust prediction of specimen movements. J. Struct. Biol., 152, 36–51.

35. Zheng, S.Q., Palovcak, E., Armache, J.-P., Verba, K.A., Cheng, Y. and Agard, D.A. (2017) MotionCor2: anisotropic correction of beam-induced motion for improved cryo-electron microscopy. Nat. Methods, 14, 331–332.

36. Rohou, A. and Grigorieff, N. (2015) CTFFIND4: Fast and accurate defocus estimation from electron micrographs. J. Struct. Biol., 192, 216–221.

37. Kucukelbir, A., Sigworth, F.J. and Tagare, H.D. (2014) Quantifying the local resolution of cryo-EM density maps. Nat. Methods, 11, 63–65.

38. Liebschner, D., Afonine, P.V., Baker, M.L., Bunkóczi, G., Chen, V.B., Croll, T.I., Hintze, B., Hung, L.W., Jain, S., McCoy, A.J., et al. (2019) Macromolecular structure determination using X-rays, neutrons and electrons: recent developments in Phenix. Acta Crystallogr D Struct Biol, 75, 861–877.

39. Pettersen, E.F., Goddard, T.D., Huang, C.C., Couch, G.S., Greenblatt, D.M., Meng, E.C. and Ferrin, T.E. (2004) UCSF Chimera--a visualization system for exploratory research and analysis. J. Comput. Chem., 25, 1605–1612.

40. Emsley, P. and Cowtan, K. (2004) Coot: model-building tools for molecular graphics. Acta Crystallogr. D Biol. Crystallogr., 60, 2126–2132.

41. Yu, C.H., Bhattacharya, A., Persaud, M., Taylor, A.B., Wang, Z., Bulnes-Ramos, A., Xu, J., Selyutina, A., Martinez-Lopez, A., Cano, K., et al. (2021) Nucleic acid binding by SAMHD1 contributes to the antiretroviral activity and is enhanced by the GpsN modification. Nat. Commun., 12, 731.

42. Alexandrov, K., Scheidig, A.J. and Goody, R.S. (2001) Fluorescence methods for monitoring interactions of Rab proteins with nucleotides, Rab escort protein, and geranylgeranyltransferase. Methods Enzymol., 329, 14–31.

43. Bai, N., Roder, H., Dickson, A. and Karanicolas, J. (2019) Isothermal Analysis of ThermoFluor Data can readily provide Quantitative Binding Affinities. Sci. Rep., 9, 2650.

44. Ji, X., Wu, Y., Yan, J., Mehrens, J., Yang, H., DeLucia, M., Hao, C., Gronenborn, A.M., Skowronski, J., Ahn, J., et al. (2013) Mechanism of allosteric activation of SAMHD1 by dGTP. Nat. Struct. Mol. Biol., 20, 1304–1309.

45. Traut, T.W. (1994) Physiological concentrations of purines and pyrimidines. Mol. Cell. Biochem., 140, 1–22.

46. Seamon, K.J., Hansen, E.C., Kadina, A.P., Kashemirov, B.A., McKenna, C.E., Bumpus, N.N. and Stivers, J.T. (2014) Small molecule inhibition of SAMHD1 dNTPase by tetramer destabilization. J. Am. Chem. Soc., 136, 9822–9825.

