## Supplemental information for "Phosphorylation of SAMHD1 Thr592 increases C-terminal domain dynamics, tetramer dissociation, and ssDNA binding kinetics"

### Phosphorylation or phosphomimetic mutation of SAMHD1 Thr592 increases C-terminal domain dynamics, GTP release and ssDNA binding kinetics

Benjamin Orris[†], Kevin W. Huynh[§], Mark Ammirati[§], Seungil Han[§], Ben Bolaños[θ], Jason Carmody[θ], Matthew D. Petroski[θ], Benedikt Bosbach[◦], David J. Shields[◦], James T. Stivers[†,*]

**Supplemental Tables**

**Table S1: Single-stranded DNA oligonucleotide sequences.**

| Oligo | Sequence |
|---|---|
| ssDNA57 | 5'-TGG AGA ATC CCG GTG CCG AGG CCG CTC AAT TGG TCG TAG ACA GCT CTA GCA CCG CTT-3' |
| ssDNA90 | 5'-GGC TCG ACA ATT GAT CCT GTA TGC CTC AGC TTA GCT ATC GAT ATA GCT ATC GAT TCC TCA GCA TTG TGT GAA TTG TGA CTA GCG GAT ACC-3' |
| psDNA5 | 5'-C*G*C*C*T-3'<br>* indicates non-bridging phosphorothioate linkage |

**Table S2: TSA-CRAFT program output.**

| Enzyme | Condition | Replicate | $T_m$ | $R^2$ | Hill Slope |
|---|---|---|---|---|---|
| Dye Alone | n/a | 1 | 58.0516 | 0.989389 | 0.683353 |
| Dye Alone | n/a | 2 | 58.1158 | 0.985737 | 0.575027 |
| Dye Alone | n/a | 3 | 57.5747 | 0.991723 | 0.842918 |
| Wild-Type | 2 mM dGTPαS | 1 | 77.0872 | 0.99653 | 0.956088 |
| Wild-Type | 2 mM dGTPαS | 2 | 77.0708 | 0.997079 | 0.892037 |
| Wild-Type | 2 mM dGTPαS | 3 | 77.3963 | 0.998571 | 0.912561 |
| T592E | 2 mM dGTPαS | 1 | 75.863 | 0.998856 | 1.21081 |
| T592E | 2 mM dGTPαS | 2 | 75.7707 | 0.998168 | 1.05329 |
| T592E | 2 mM dGTPαS | 3 | 76.299 | 0.996948 | 1.25929 |
| pSAMHD1 | 2 mM dGTPαS | 1 | 75.7236 | 0.99811 | 0.982312 |
| pSAMHD1 | 2 mM dGTPαS | 2 | 75.7859 | 0.99792 | 0.988214 |
| pSAMHD1 | 2 mM dGTPαS | 3 | 76.5148 | 0.998262 | 1.05413 |
| Δ583-626 | 2 mM dGTPαS | 1 | 74.0477 | 0.998465 | 0.898611 |
| Δ583-626 | 2 mM dGTPαS | 2 | 74.1603 | 0.998682 | 0.895201 |
| Δ583-626 | 2 mM dGTPαS | 3 | 74.6798 | 0.99844 | 0.897216 |
| Δ600-626 | 2 mM dGTPαS | 1 | 77.5901 | 0.999491 | 1.02777 |
| Δ600-626 | 2 mM dGTPαS | 2 | 77.6032 | 0.999409 | 1.03033 |
| Δ600-626 | 2 mM dGTPαS | 3 | 77.7691 | 0.999394 | 1.03357 |
| Wild-Type | Enzyme alone | 1 | 44.2739 | 0.997981 | 1.477 |
| Wild-Type | Enzyme alone | 2 | 44.3784 | 0.998731 | 1.41723 |
| Wild-Type | Enzyme alone | 3 | 44.4327 | 0.997741 | 1.43064 |
| T592E | Enzyme alone | 1 | 44.5379 | 0.998901 | 1.46469 |
| T592E | Enzyme alone | 2 | 44.6177 | 0.998745 | 1.39772 |
| T592E | Enzyme alone | 3 | 44.7762 | 0.998302 | 1.41055 |
| pSAMHD1 | Enzyme alone | 1 | 45.5318 | 0.999189 | 1.47642 |
| pSAMHD1 | Enzyme alone | 2 | 45.5427 | 0.998815 | 1.46445 |
| pSAMHD1 | Enzyme alone | 3 | 45.5842 | 0.99904 | 1.41357 |
| Δ583-626 | Enzyme alone | 1 | 45.1876 | 0.999028 | 1.47663 |
| Δ583-626 | Enzyme alone | 2 | 45.2476 | 0.999102 | 1.45138 |
| Δ583-626 | Enzyme alone | 3 | 45.2119 | 0.998564 | 1.46097 |
| Δ600-626 | Enzyme alone | 1 | 45.9235 | 0.998834 | 2.00678 |
| Δ600-626 | Enzyme alone | 2 | 46.0033 | 0.998562 | 2.13095 |
| Δ600-626 | Enzyme alone | 3 | 45.9238 | 0.998504 | 2.00854 |

**Table S3: Cryo-EM data collection, processing, and refinement statistics[1]**

| | |
|---|---|
| **Data Collection and Processing** | |
| Magnification | 130,000x |
| Voltage (kV) | 300 |
| Total Electron Exposure ($e^-/Å^2$) | 57.4 |
| Defocus Range (µm) | -0.6 to -3.2 |
| Pixel Size (Å) (super resolution mode) | 0.420 |
| Movies Recorded | 3,222 |
| Symmetry Imposed | D2 |
| Initial particle images (no.) | 1,030,155 |
| Final particle images (no.) | 114,078 |
| Map Resolution at FSC=0.143 (Å) | 2.89 |
| **Refinement** | |
| Map sharpening *B* factor ($Å^2$) | -71.75 |
| Model composition in the asymmetric unit: | |
| Non-hydrogen atoms | 16,272 |
| Protein residues | 1,948 |
| Average B factors ($Å^2$) | |
| All atoms | 65.0 |
| R.M.S. Deviations | |
| Bond lengths (Å) | 0.003 |
| Bond angles (°) | 0.508 |
| Validation | |
| MolProbity score | 1.64 |
| Clashscore | 7.92 |
| Poor rotamer (%) | 0.0 |
| Ramachandran Plot | |
| Favored (%) | 96.70 |
| Allowed (%) | 3.30 |
| Outliers (%) | 0.00 |

[1]EMDB: EMD-26567, PDB: 7UJN

**Table S4: Kinetic analysis results with confidence intervals.**

| | Wild-Type | | T592E | |
|---|---|---|---|---|
| **Parameter** | **Value** | **95% confidence interval** | **Value** | **95% confidence interval** |
| $k_{cat}^{dTTP}$ | 4.6 $s^{-1}$ | 3.3 to 8.8 $s^{-1}$ | 3.6 $s^{-1}$ | 3.0 to 4.5 $s^{-1}$ |
| $K_m^{dTTP}$ | 20.1 mM | 16.7 to 27.6 mM | 12.8 mM | 10.1 to 17.4 mM |
| $K_{act}^{GTP}$ | 0.17 mM | 0.01 to 0.06 mM | 0.42 mM | 0.01 to 0.08 mM |
| $\alpha$[a] | 7 | 5 to 10 | | |

[a]See eq 1-4 in methods. This unitless parameter, which reflects the fold decrease in $K_m^{dTTP}$ when GTP is saturating, was not discernably different for the wild-type enzyme and T592E. A shared value was used in the fitting.

#### SUPPLEMENTAL METHODS

##### Course-grained normal mode analysis R script:

```
library(bio3d)
#Read tetrameric SAMHD1 structure 6TXC & pull monomer from it
SAMHD1.full <- read.pdb("6txc")
SAMHD1.tet <- trim.pdb(SAMHD1.full, atom.select(SAMHD1.full, chain="A"))

#Normal Mode Analysis
modes <- nma(SAMHD1.tet)
#Plot results
plot(modes, sse = SAMHD1.tet)

#Export pymol movie and vector representation of trajectories
mktrj(modes, mode=7)
pymol(modes, mode=7)

#Determine cross-correlation matrix and render in pymol
cm <- dccm(modes)
pymol(cm, SAMHD1.tet, type="launch", exefile = "C:/Users/bporr/PyMOL/PyMOLWin.exe")

#Calculate deformation energies and fluctuations
defe <- deformation.nma(modes)
defsums <- rowSums(defe$ei[,1:3])
flucts <- fluct.nma(modes, mode.inds=seq(7,9))

#Render deformation energies and fluctuations in pymol
write.pdb(pdb=NULL, xyz=modes$xyz, file="SAMHD1-defor.pdb", b=defsums)
write.pdb(pdb=NULL, xyz=modes$xyz, file="SAMHD1-fluct.pdb", b=flucts)
```

#### SUPPLEMENTAL FIGURES

**A**

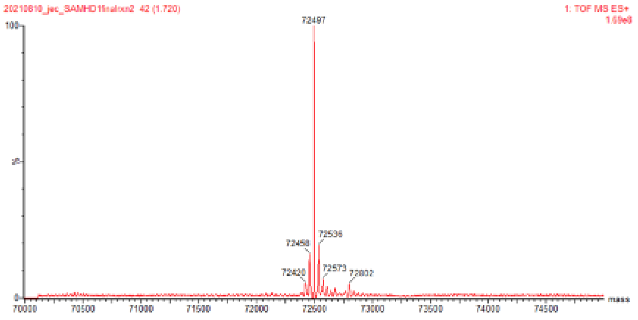

**B**

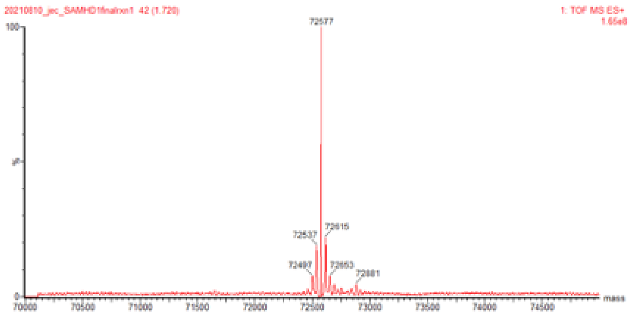

**C** **SAMHD1 97% sequence coverage (425 peptides)**

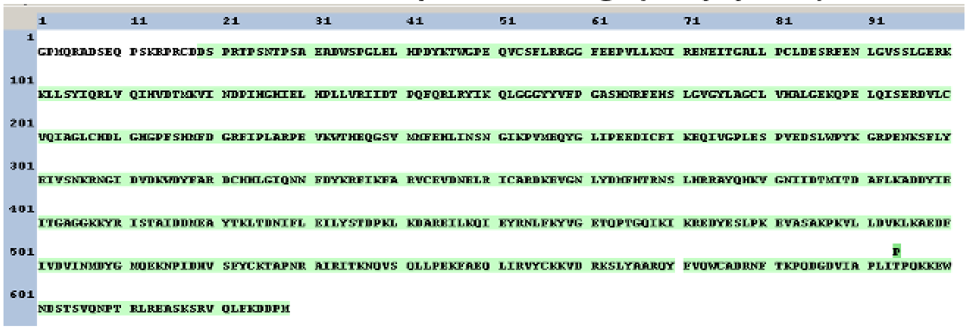

**HCD Peptide Fragmentation Assignment** **Phospho-T592**

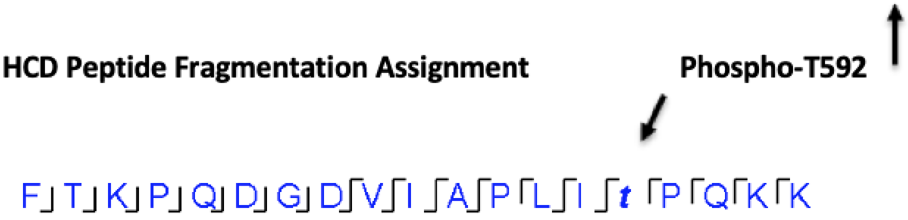

**Figure S1: Characterization of SAMHD1 phosphorylated by CDK2/cyclin E1 by mass spectrometry.** (A) Mass spectrum of unreacted SAMHD1 (B) Mass spectrum of SAMHD1 phosphorylated by CDK2/cyclin E1 in the presence of 375 μM ATP for 1 h. (C) Inline digest and LC-MS2 analysis provided 97% sequence coverage for reacted SAMHD1, with only T592 identified as the site of phosphorylation on all four T592 containing peptides. MS2 fragmentation (HCD) of these peptides consistently assigned T592 as the site of phosphorylation.

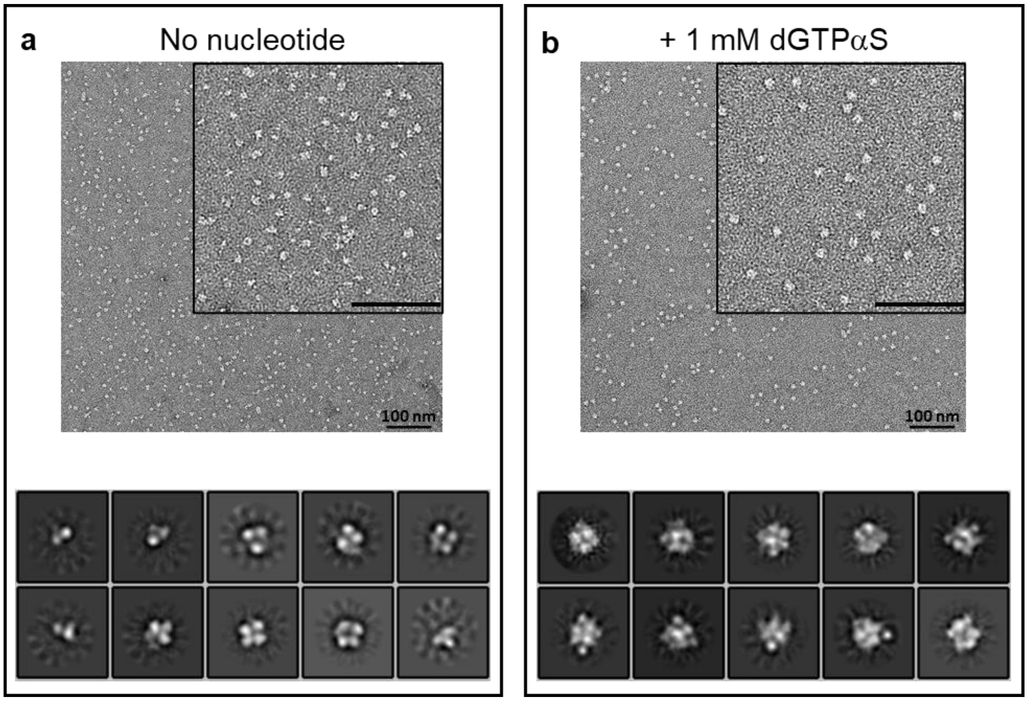

**Figure S2. Negative stain EM of SAMHD1-dGTPαS complex tetramer.** (**A**) Without nucleotide, heterogenous particles are observed in raw micrographs (panel a, top). 2D class averages (panel a, bottom) reveal a mixture of classes with particles that differ in size. (**B**) After nucleotide addition, homogeneous particles are observed (panel b top). 2D class averages (panel b, bottom) reveal particles that increase in volume and are consistent with tetramer formation.

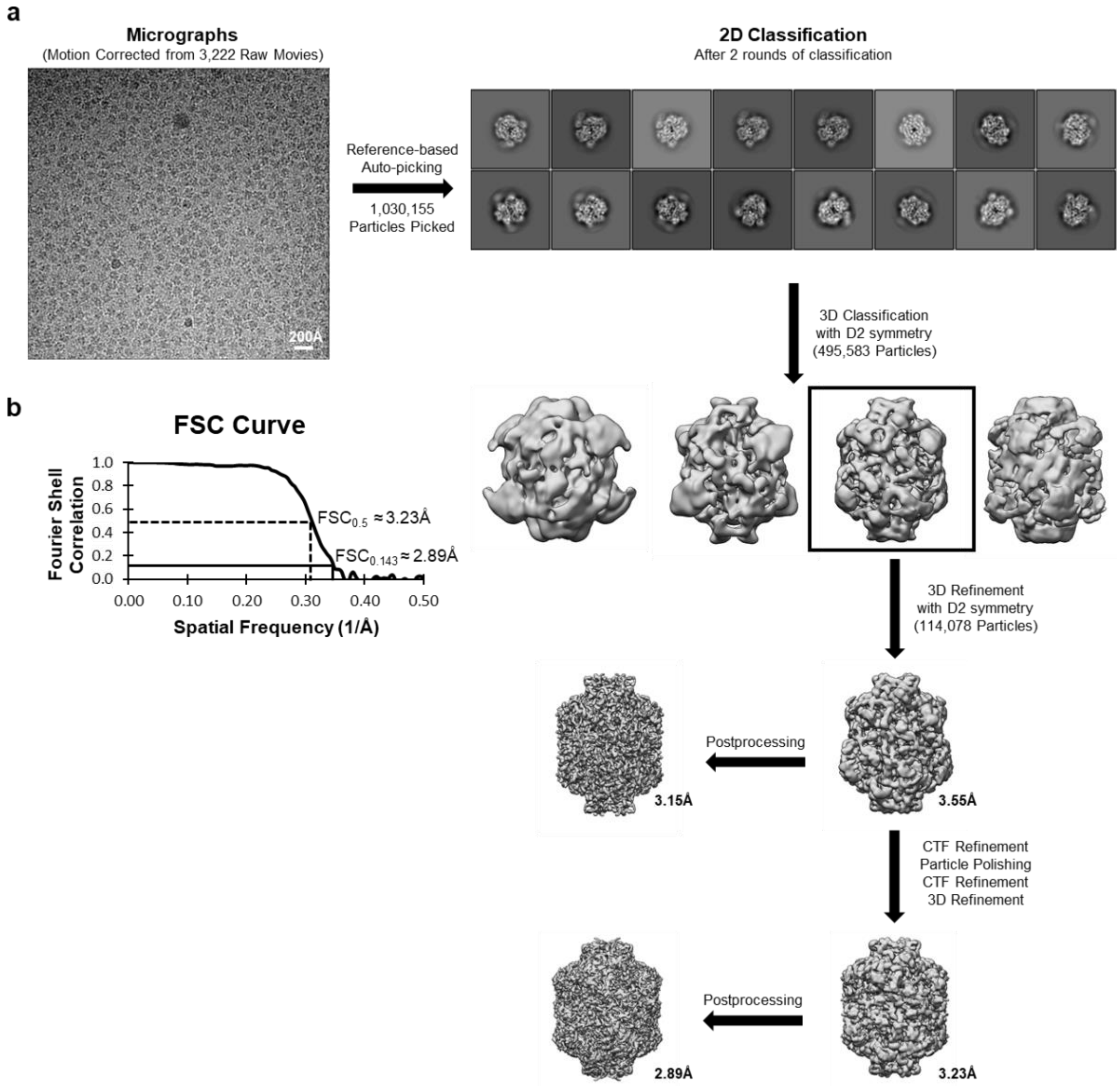

**Figure S3.** General cryo-EM data processing workflow for the full-length hSAMHD1 with dGTP$\alpha$S. **b)** FSC curve for the final 2.98Å cryo-EM map.

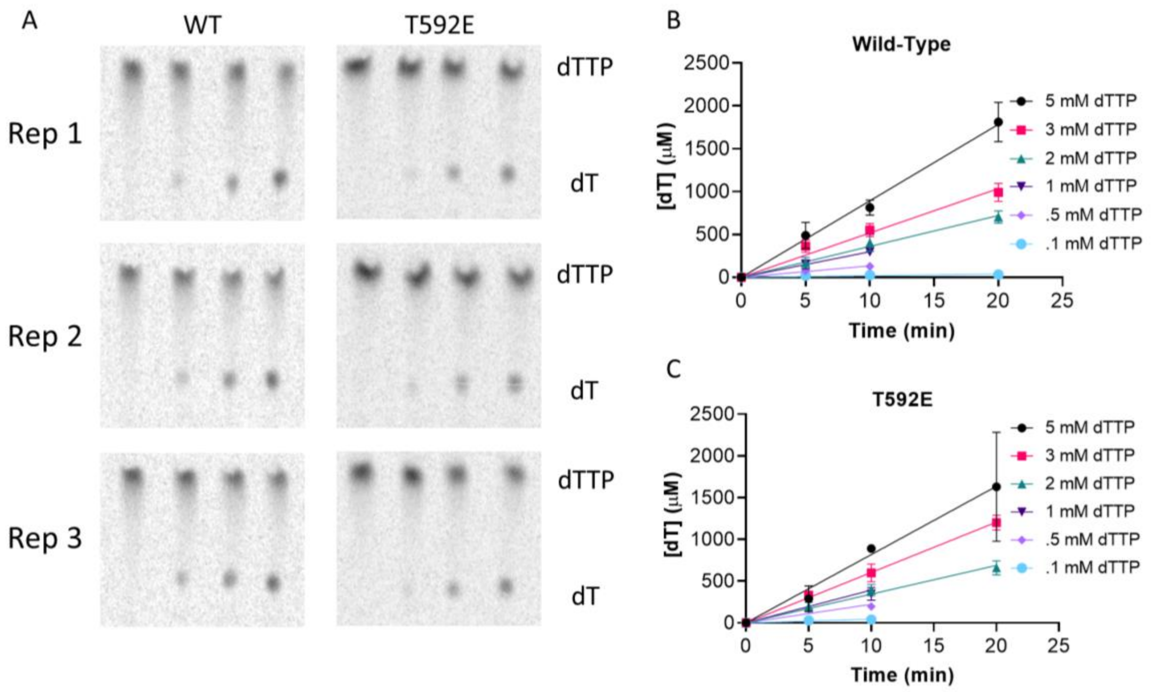

**Figure S4. Kinetic analysis method verification. (A)** Reversed phase TLC chromatograms of the hydrolysis of 5 mM dTTP by SAMHD1 (0.5 µM) in the presence of 0.5 mM GTP. Fractions were withdrawn at 0, 5, and 10, and 20 minutes. **(B)** Linear regression [dT] formation vs time at varying concentrations of dTTP by SAMHD1 (0.5 µM) in the presence of 0.5 mM GTP. Error bars indicate standard error of mean from three replicate reactions.

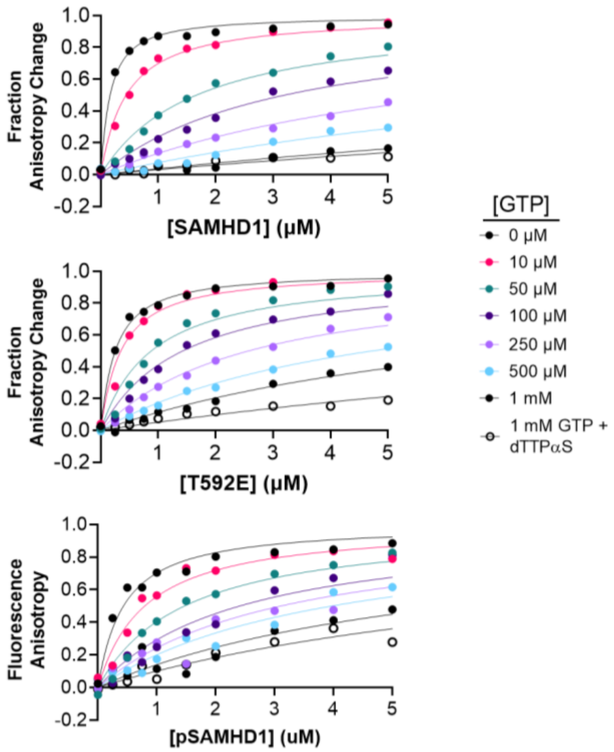

**Figure S5. GTP competition with psDNA5 binding.** Binding of SAMHD1 to 5'FAM labeled psDNA5 in the presence of increasing concentrations of GTP or in the presence of a combination of GTP and dTTPαS.

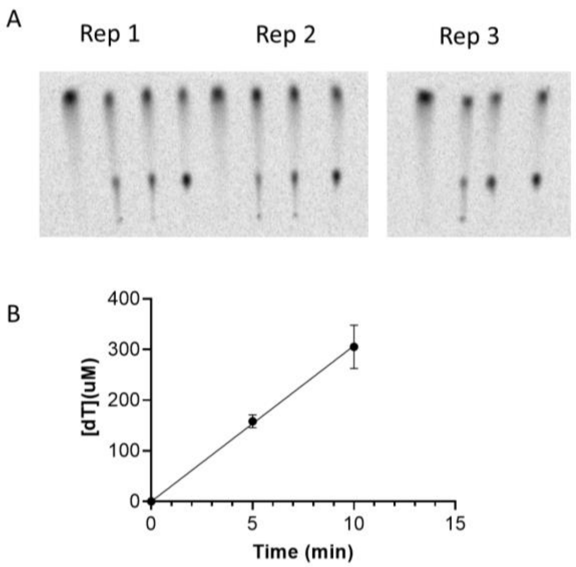

**Figure S6. Mant-GTP activation of SAMHD1 dNTPase. (A)** Reversed-phase TLC chromatograms of the hydrolysis of 1 mM dTTP by SAMHD1 (0.5 µM) in the presence of 1 mM mant-GTP. **(B)** Linear regression of [dT] formation vs time to determine initial reaction rate. Error bars indicate standard error of mean from thee replicate reactions.

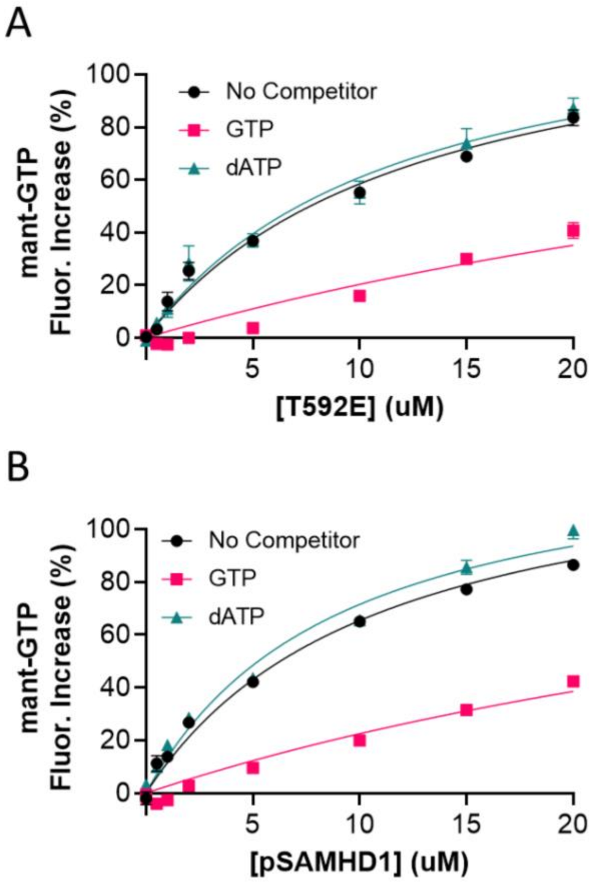

**Figure S7. Binding of mant-GTP to SAMDH1 T592E and pSAMHD1 (uncorrected for non-specific binding). (A)** Binding of SAMHD1 T592E to mant-GTP (0.5 µM) in the presence of dATP (1 mM) or GTP (1 mM). **(B)** Binding of pSAMHD1 to mant-GTP (0.5 µM) in the presence of dATP (1 mM) or GTP (1 mM). Error bars indicate standard error of mean determined by three replicate titrations.

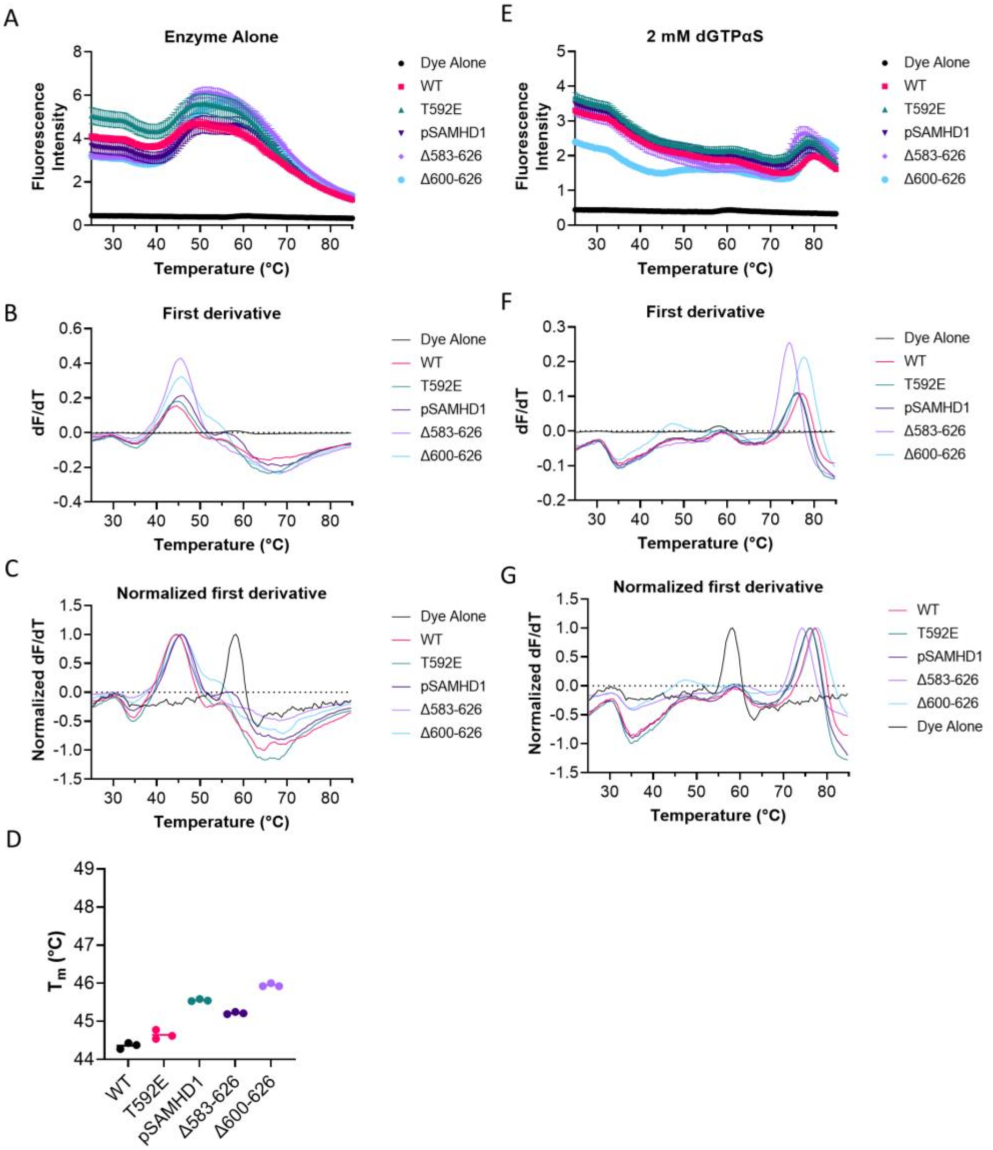

**Figure S8. TSA full data analysis. (A)** Averaged thermal melt raw data in the absence of dGTPαS. Error bars indicate standard error of mean from three replicates. **(B)** First derivative plot of (A). **(C)** Normalization of (B). **(D)** Melt temperatures calculated for individual replicates of each enzyme in TSA-CRAFT. **(E)** Averaged thermal melt raw data in the presence of 2 mM dGTPαS. Error bars indicate standard error of mean from three replicates. **(F)** First derivative plot of (E). **(G)** Normalization of (F).

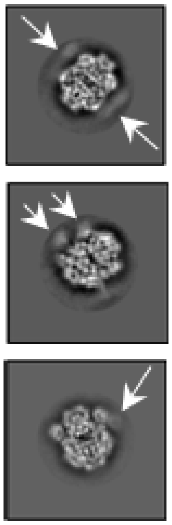

**Figure S9.** Cryo-EM 2D class averages with the arrows highlighting the hazy densities that are likely the flexible SAM domains.

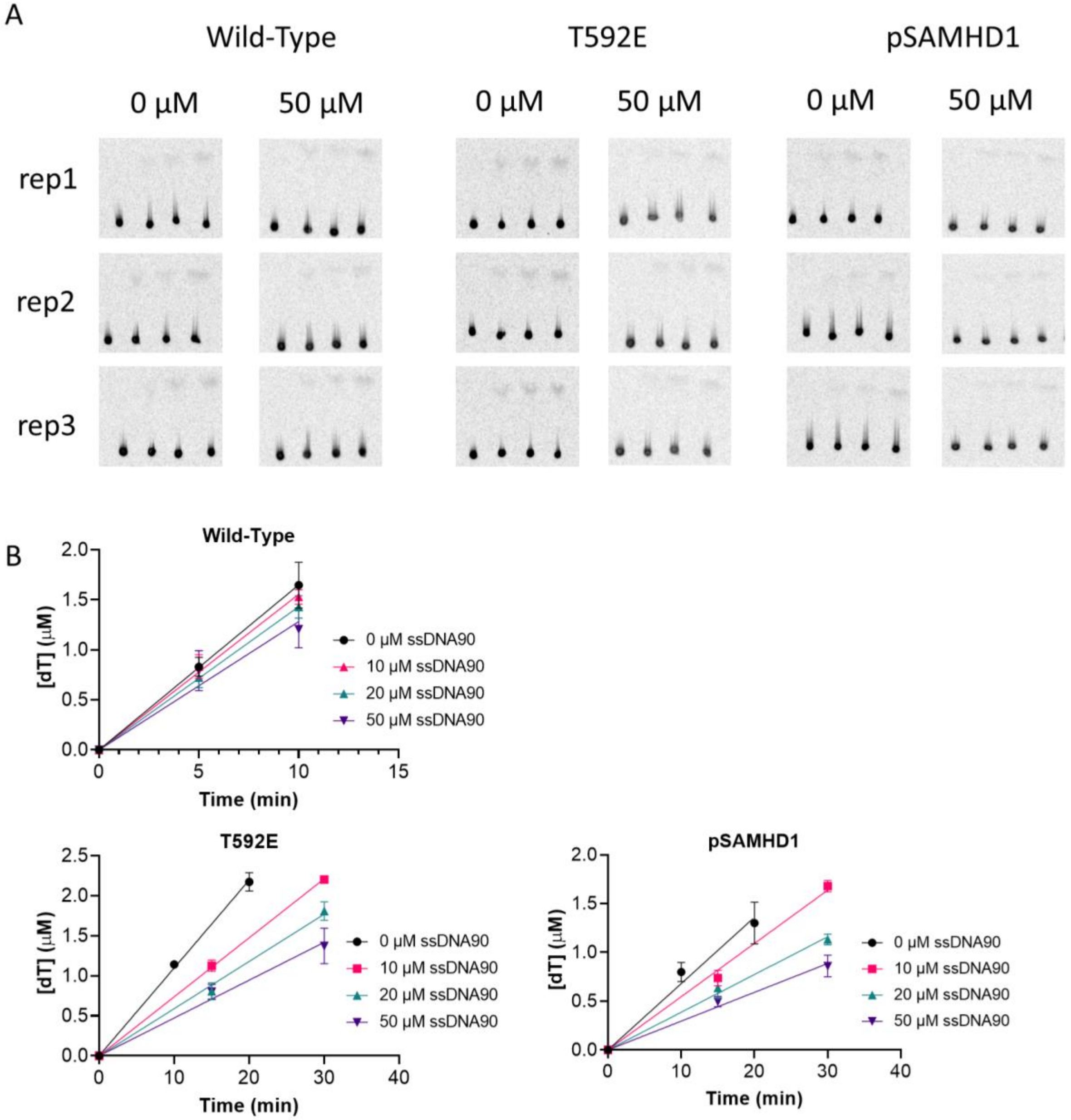

**Figure S10. ssDNA competition with SAMHD1 dNTPase. (A)** PEI-Cellulose TLC chromatograms of dTTP (10 µM) hydrolysis by SAMHD1 (0.5 µM) in the presence of 10 µM GTP and either 0 or 50 µM ssDNA90. **(B)** Linear regression of initial reaction rates from (A). Error bars indicate standard error of mean from three replicate reactions.
